# Efficient capture–recapture inference for spatially varying natal dispersal, survival and recruitment

**DOI:** 10.64898/2026.08.28.747721

**Authors:** Matia H. Muller, Fabian R. Ketwaroo, Wolfgang Fiedler, Olaf Geiter, Christof Herrmann, Michael Schaub

## Abstract

1. Natal dispersal is a key process in population ecology because it links local demographic processes to broader-scale population dynamics by redistributing individuals. When using capture–recapture data, multistate capture–recapture models using discrete spatial units as states are the gold standard for estimating natal dispersal among spatial units while accounting for spatial variation in survival, recruitment and imperfect detection. However, because their computational cost increases rapidly with the number of spatial units, applications have been limited to a small number of units. Therefore, in practice, these models cannot provide spatially detailed inference on natal dispersal across large landscapes.
2. We develop a computationally efficient Bayesian capture-recapture model, called the efficient natal dispersal (END) model, to estimate natal dispersal among discrete spatial units jointly with spatial variation in demographic parameters and detection probabilities. The END model relies on two key structural features: juveniles and breeders are separated into two arrays, and resightings outside the natal spatial unit are aggregated over time for individuals released as juveniles.
3. Using simulations, we show that the END model is considerably (up to 30 times) more computationally efficient than a conventional multistate model, while maintaining comparable parameter accuracy. We then apply the END model to white stork (*Ciconia ciconia*) capture-recapture data from Germany across 101 hexagonal spatial units, a spatial resolution at which a conventional multistate model is computationally infeasible. We estimate natal dispersal among units jointly with spatial variation in survival and recruitment. This allows us to identify areas of lower or higher survival, earlier or delayed recruitment, and dispersal probabilities among all units. By combining estimated dispersal probabilities with existing data on the number of juveniles born in each spatial unit, we estimate natal dispersal in terms of numbers of individuals and identify units with positive or negative net migration, sources and sinks.
4. Overall, our approach moves capture–recapture analyses from estimating natal dispersal among a few spatial units to inferring dispersal networks and assessing their demographic consequences across large domains. Our approach is applicable to many spatially structured capture–recapture datasets, opening new opportunities for studying spatial population dynamics.

## Introduction

Understanding the spatial structure of population dynamics is a central goal in population ecology (Gurevitch et al., 2016). This structure is shaped by spatial variation in demographic parameters such as survival and recruitment (e.g., Muller et al., 2026), but also by the dispersal of individuals, which redistributes individuals across space and thereby influences local population growth (Johst & Brandl, 1997). Developing methods to estimate dispersal is therefore essential to understand population dynamics across space. This is particularly true for dispersal between birth and first breeding, i.e., natal dispersal, which is generally more frequent and occurring over larger distances compared to dispersal between breeding attempts (Greenwood & Harvey, 1982; Harts et al., 2016), and is therefore expected to play a more important role in shaping spatial population dynamics than breeding dispersal.

To study natal dispersal across large areas, the most widely available data are capture– recapture data, in which individuals are marked at one location and may later be re-encountered either at the same location or elsewhere. To obtain dispersal estimates from such data, multistate capture-recapture models that use discrete spatial units as states are commonly used (Arnason, 1973; Lebreton et al., 2003). These models can estimate natal dispersal probability from one spatial unit to another while accounting for imperfect detection and spatial variation in survival, recruitment and detection probabilities. Thus, they are the gold standard to estimate dispersal of individuals across discrete spatial units. When combined with additional demographic data, the dispersal estimates these models can then be used for estimating how many individuals are expected to move from one spatial unit to another, whether a given unit acts as a net importer or exporter of individuals, or whether it functions as a source or a sink (Pulliam, 1988; Runge et al., 2006).

Multistate models become computationally expensive when applied to large numbers of individuals. This limitation can be overcome by using the multinomial formulation, which uses aggregated data and is therefore unaffected by the number of individuals (Schaub & Kéry, 2022). However, even with a multinomial formulation, conventional multistate models become computationally demanding when the number of spatial units increases (Lagrange et al., 2014). The problem is further exacerbated when the model includes further complexity, for example by accounting for temporal variability in different demographic parameters, by considering different age classes, or by accounting for different conditions for individuals at detection (e.g., breeder, non-breeder). This computational bottleneck prevents conventional multistate models from reconstructing spatially detailed dispersal networks across large landscapes, motivating the development of models that retain the main inferential advantages of multistate models at reduced computational cost.

Here, we develop a novel model, the efficient natal dispersal (END) model, to estimate natal dispersal among discrete spatial units jointly with unit-specific survival and recruitment, while reducing the computational cost relative to a conventional multistate model. Cost reduction comes from using separate arrays for juveniles and breeders and removing the temporal dimension from resightings outside the natal spatial unit for individuals released as juveniles. We compare estimation accuracy and computational efficiency of the END model against a conventional multistate model using several simulation scenarios. We then apply the END model to a dataset of white storks in Germany containing 88,445 individuals across a network of 101 spatial units, for which a conventional multistate model is computationally infeasible. To demonstrate the potential of our new model, we use the estimated dispersal network between spatial units together with data on the local number of fledglings produced to calculate the expected number of juveniles moving among spatial units. This allows us to quantify immigration and emigration patterns and identify source and sink units.

## Methods

### Conventional multistate capture-recapture model

The conventional multistate capture–recapture model follows the framework described by Lebreton et al. (2003). Across several spatial units, it estimates three demographic processes: survival, recruitment into the breeding population, and dispersal among spatial units, as well as an observation process: detection probability. Survival, recruitment and detection probability are allowed to vary among spatial units and age classes. Once recruited, individuals are assumed to remain breeders until death. Non-breeders are not detectable. In contrast to Lebreton et al. (2003), however, we consider natal dispersal only; breeding dispersal does not occur in our framework.

In the conventional multistate model, individuals are described by a set of discrete states, which typically correspond to a combination of spatial unit, age class and status (breeder or non-breeder). These states represent the latent biological states of individuals, some of which are observable, and define the structure of the model. Here, we implement this model using the multinomial formulation, because of its efficient computation compared to the state-space formulation (Schaub & Kéry 2022). The individual capture histories are transformed into a multistate m-array **m** (Burnham, 1987). Each row of **m** corresponds to a state-time combination at which individuals are released, and each column corresponds to the state-time combination of their first subsequent re-encounter, or to the event of never being re-encountered (last column). Each cell of **m** contains therefore the number of individuals observed in a given state-time combination and first re-encountered in another state-time combination, or never re-encountered. Supplement: Figure S1 shows how individual capture-histories are aggregated into an m-array.

Transition probabilities describe how individuals move among states between two consecutive years, as a function of the demographic parameters: survival, recruitment and dispersal among spatial units. In the multinomial formulation, these transition probabilities are combined with state-specific detection probabilities to calculate the probabilities ***π*** corresponding to the cells of **m**. For each state-time combination at which individuals are detected, the corresponding row of **m** follows a multinomial distribution with cell probabilities given by the corresponding row of ***π***. Specifically, for each row **m**_*K*_ of the m-array, then

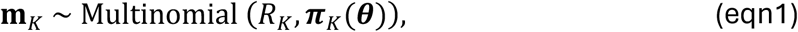

where *R*_*K*_ is the total number of individuals released in state-time combination *K*, and ***π***_*K*_(***θ***) is the corresponding vector of probabilities defined by the matrix ***π***, which is a function of the underlying demographic and observation parameters ***θ*** (Supplement: Table S1 for symbol definitions). The full likelihood of the model is obtained by combining the multinomial contributions over all rows of **m**:

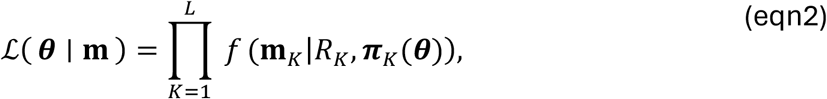

where *f* denotes the multinomial probability mass function and *L* is the number of rows in **m**.

### Efficient natal dispersal (END) model

The computational advantage of the END model over the conventional multistate model relies on two modifications: (i) treating juveniles, which may disperse to any spatial unit, separately from breeders, which are assumed to no longer disperse, and (ii) aggregating across years resightings outside the natal spatial unit for individuals released as juveniles. Like the conventional model, the END model estimates parameters using a multinomial likelihood, but organizes the capture–recapture data into two modified m-arrays, termed the juvenile-END-array and the breeder-END-array. Their structures differ from the conventional multistate m-array, and their cell probabilities are defined differently.

In the juvenile-END-array **m**^*juv*^, rows represent juveniles released in a given natal spatial unit and year, ordered first by natal spatial unit and then by release year. Let *T* denote the number of years and *S* the number of spatial units. For each row, the first *T* − 1 columns correspond to the first re-encounter in the natal spatial unit, with one column for each year (starting at year 2, as no re-encounter is possible the first year). The following *S* columns correspond to the first re-encounter in each possible destination spatial unit, irrespective of the year in which this re-encounter occurred. The destination column corresponding to the natal spatial unit is structurally zero, because re-encounters in the natal spatial unit are already represented by the year-specific columns. The final column corresponds to individuals never re-encountered. The dimension of the juvenile-END-array is (*T* − 1)*S* × (*T + S*), which is substantially smaller than the m-array of the conventional model (*T* − 1)*S* × ((*T* − 1)*S +* 1).

The breeder-END array **m**^*breed*^ is used for individuals detected after the juvenile stage, i.e., breeders that are assumed to not disperse anymore. Its rows are ordered first by spatial unit, then by release year, and finally by breeder age class. The columns represent the first re-encounter in the same spatial unit, with one column for each year, followed by a final column for individuals never encountered again. Let *A* denote the number of breeder age classes. The dimension of **m**^*breed*^ is (*T* − 1)*SA* × *T*. Supplement: Figure S2 shows how juvenile- and breeder-END-arrays are constructed from individual capture-histories.

The cell probabilities of the two END-arrays are computed in several steps. First, we compute the probability *q*_*s,t,k*_ that a juvenile that has dispersed to spatial unit *s* from year *t* to year *t +* 1 and is available for detection (i.e., has recruited) from year *t +* 1 onward, is first detected in year *k*:

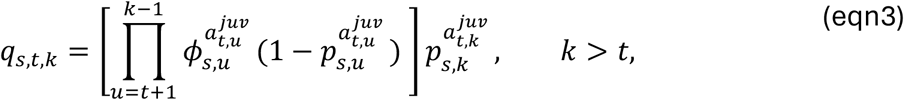

where 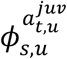 is the survival in spatial unit *s* in year *u* for age class 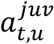 (i.e., the age class in year *u* of an individual released as a juvenile in year *t*) and 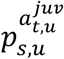 the probability that a breeder is detected at age class 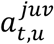 in spatial unit *s* and year *u*.

Next, we compute the probability that an individual that dispersed to spatial unit *s* from year *t* to year *t +* 1 is first detected in year *k*, after accounting for delayed recruitment (*h*_*s,t,k*_). In species where recruitment is delayed, we define

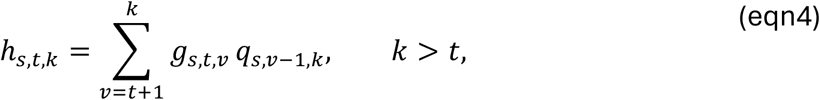

where *g*_*s,t,v*_ is the probability that an individual that dispersed to spatial unit *s* from year *t* to year *t +* 1 survives and recruits in year *v*, thereby becoming available for detection from year *v* onward. When recruitment occurs immediately after natal dispersal such as in short-lived species, *g*_*s,t,t* +1_ = 1, and therefore *h*_*s,t,k*_ = *q*_*s,t,k*_. Otherwise, *g*_*s,t,v*_ is computed from recruitment and survival probabilities (Appendix S1).

We then use *h*_*s,t,k*_ to compute the cell probabilities ***π***^*juv*^(***θ***) of the juvenile-END-array. For juveniles released in natal spatial unit *s* and year *t*, the probability of being first re-encountered in the natal spatial unit in year *k*, corresponding to column *k* − 1, is

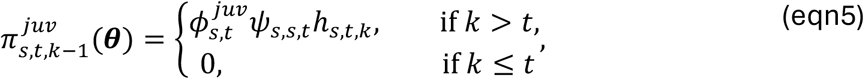

where 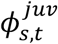 is juvenile survival in spatial unit *s* at time *t* and *ψ*_*s,s,t*_ is the probability of remaining in spatial unit *s* at time *t* (hereafter, fidelity).

The probability of being first re-encountered, irrespective of the year, in any spatial unit *r*, corresponding to column (*T* − 1) *+ r*, is given by

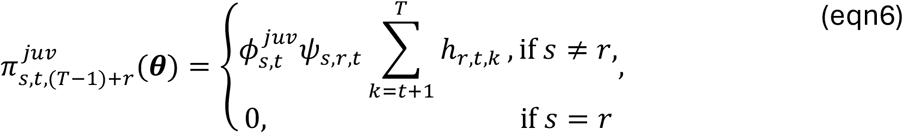

where *ψ*_*s,r,t*_ is the probability of natal dispersal from spatial unit *s* to spatial unit *r* at time *t*, and 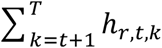 is the probability of being detected at least once in spatial unit *r* after having dispersed to spatial unit *r* at time *t*.

The probability of never being re-encountered in any spatial unit, corresponding to column *T + S*, is computed as the complement of one:

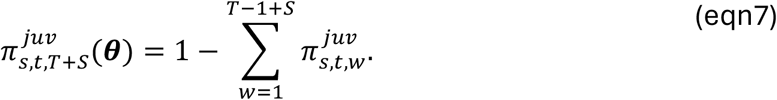

As each row 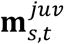 of the juvenile-END array corresponds to juveniles released in a given natal spatial unit *s* and year *t*, the likelihood contribution of this row is then

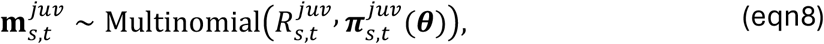

where 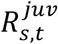 is the number of juveniles released in the spatial unit *s* at time *t*, and 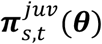 is the vector of cell probabilities defined above.

We next compute the cell probabilities and the likelihood of the breeder-END-array. For individuals released in spatial unit *s*, year *t*, and breeder age class *a*_*rel*_, the probability of being first re-encountered in the same spatial unit in year *k*, corresponding to column *k* − 1, is

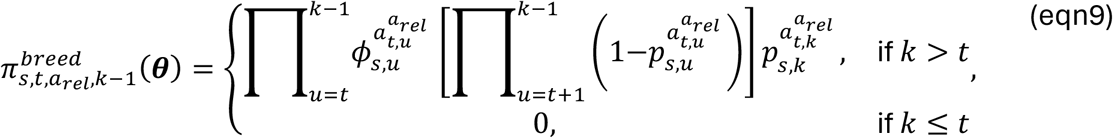

where 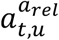 is the age class in year *u* of an individual released in age class *a*_*rel*_ in year *t*.

If survival and detection do not vary with age among breeders, as commonly assumed, the precomputed *q*_*s,t,k*_ quantities can also be used to simplify the computation of the breeder-END cell probabilities:

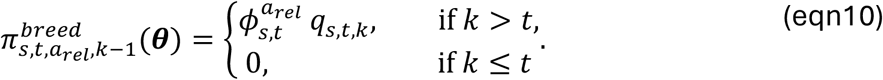

This reduces the number of arithmetic operations because the post-recruitment detection process does not need to be recomputed separately for each breeder-END-array row.

Finally, each row 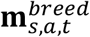 of the breeder-END-array follows a multinomial distribution,

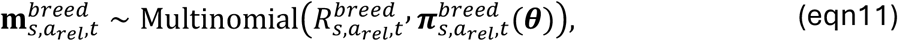

where 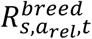 is the number of breeders released in spatial unit *s*, in age class *a*_*rel*_ and at time *t*, and 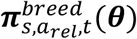 is the vector of cell probabilities defined above.

The full likelihood of the END model can therefore be expressed as:

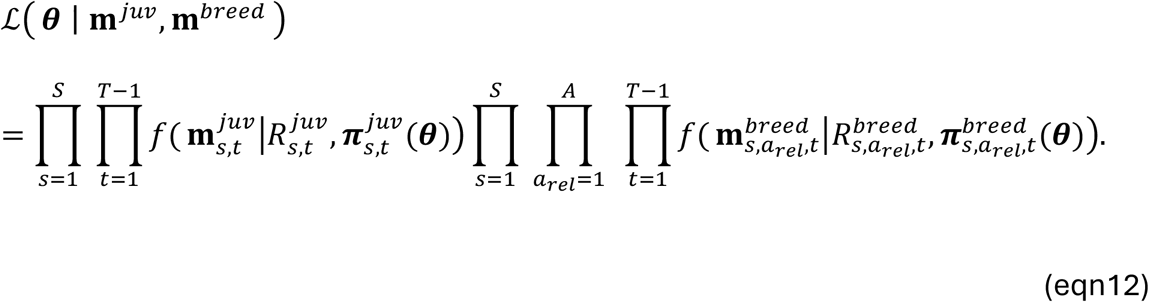

### Simulation scenarios

We considered ten different simulation scenarios (a-j) to test the performance of the END model against the conventional multistate model. All scenarios assumed a study duration of 10 years and were replicated 100 times. For each replicate, values of age-dependent survival, detection probability and natal dispersal among spatial units were drawn from predefined distributions (Supplement: Table S2) and used to simulate a capture–recapture dataset.

The first eight scenarios (a-h) used a simple demographic structure with two survival age classes, juvenile and adult, recruitment fixed at one year of age and no temporal variability. To assess how model accuracy and computational efficiency varied across data-generating conditions, they differed in 1) the number of released individuals (5 (low) or 100 (high) per year, spatial unit and age class), 2) the number of spatial units (4 or 10), and 3) the process used to simulate dispersal probabilities. In four scenarios (a-d), dispersal probabilities were simulated from a Dirichlet distribution (Kotz et al., 2001), with parameters chosen to produce on average a fidelity probability of 0.5, with some variation between simulation replicates (details in Supplement: Table S2). In the other four scenarios (e-h), natal dispersal probability *ψ*_*i,j*_ was calculated using an exponential dispersal kernel as a function of distance between spatial units:

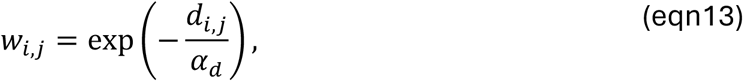

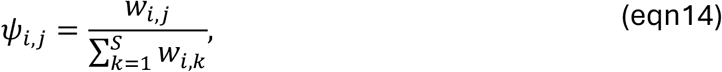

where *d*_*i,j*_ is the Euclidean distance between the centroid of spatial units *i* and *j* (Supplement: Table S3 for symbol definitions), *α*_*d*_ controls the effect of distance on dispersal, and *S* is the number of spatial units. Consequently, dispersal probabilities decrease with distance, making them more biologically realistic than the Dirichlet formulation assuming the same dispersal probabilities across all spatial units. In scenarios with four spatial units, spatial units were arranged linearly (Supplement: Fig. S3a), whereas in scenarios with 10 spatial units, they were arranged in two rows of five (Supplement: Fig. S3b), with a distance of 1 between neighbouring units.

In addition, two scenarios (i-j) were used to assess the accuracy and computational efficiency of the multistate and END models under more complex conditions. In scenario i, we included temporal variability as a fixed effect in all model parameters (survival, dispersal and detection probability). In scenario j, we considered three age classes (juvenile, 1-year-old, and adult) instead of two, with survival varying among age classes. Recruitment was allowed to occur either at 1 or 2 years of age, rather than being fixed at 1 year of age as in the other scenarios. Recruitment was drawn from a predefined distribution (Supplement: Table S2) in each replicate.

The list of scenarios is provided in Table 1.

**TABLE 1.** Simulation scenarios used to compare the END model with the conventional multistate model. The number of juveniles and adults marked refers to the number of individuals marked each year in each spatial unit.

| **Scenario** | **Spatial units** | **Juveniles marked** | **Adults marked** | **Dispersal** | **Temporal variability** | **Age classes** |
| --- | --- | --- | --- | --- | --- | --- |
| **a** | 4 | 100 | 100 | Dirichlet | No | 2 |
| **b** | 10 | 100 | 100 | Dirichlet | No | 2 |
| **c** | 4 | 5 | 5 | Dirichlet | No | 2 |
| **d** | 10 | 5 | 5 | Dirichlet | No | 2 |
| **e** | 4 | 100 | 100 | Distance | No | 2 |
| **f** | 10 | 100 | 100 | Distance | No | 2 |
| **g** | 4 | 5 | 5 | Distance | No | 2 |
| **h** | 10 | 5 | 5 | Distance | No | 2 |
| **i** | 4 | 100 | 100 | Dirichlet | Yes | 2 |
| **j** | 4 | 100 | 100 | Dirichlet | No | 3 |

### Model fitting to simulated datasets and performance assessment

All models were fitted in a Bayesian framework using Markov chain Monte Carlo (MCMC) methods in NIMBLE (de Valpine et al., 2017). We ran each model for 20,000 MCMC iterations in each of four chains, with a burn-in of 5,000 iterations and a thinning interval of 10 iterations.

For each replicate, the same simulated dataset was used for the multistate and the END model, but structured according to the requirements of each model (conventional m-array for the multistate and the two END-arrays for the END). In both models, we used uniform priors defined over the interval [0,1] for survival and detection probabilities, and for recruitment probabilities in the supplementary scenario including delayed recruitment. In scenarios where dispersal was simulated using a Dirichlet distribution, natal dispersal was assigned a symmetric Dirichlet prior in the models with all concentration parameters set to 1. In scenarios where dispersal was simulated based on distance among spatial units, natal dispersal was parameterized as in the data-generating process, i.e., as a function of distance for which a vague prior was assigned to *α*_*d*_ (uniform over [0,20]).

For each scenario and each simulation replicate, we calculated the bias as the difference between the posterior means and true values (simulated) for each parameter of interest (survival, natal dispersal, detection, recruitment, *α*_*d*_). We also computed the root mean squared error (RMSE) and the 95% credible interval (CrI) coverage of these parameters (Appendix S1).

Finally, for each simulation replicate, each scenario and each model, we quantified computational efficiency as the minimum effective sample size (ESS) across the parameters of interest divided by the computation time taken for running the four chains (MCMC efficiency; e.g., Monnahan et al., 2019).

### Case study

The White Stork (*Ciconia ciconia*) is a long-lived bird species in which natal dispersal can occur over long distances, whereas breeding dispersal is rare (Chernetsov et al., 2006). Between 2000 and 2023, 88,445 stork nestlings were marked throughout Germany, some of which were later resighted as breeder. To study spatial variation in demographic parameters and natal dispersal, we divided Germany (total surface: 357,592 km^2^) into a grid of 101 hexagonal spatial units of 7,600 km^2^ each (distance between the centroids of adjacent units: 93.7 km), some of which only partially overlapped the country (Supplement: Fig. S15). The large number of spatial units makes the conventional multistate model computationally infeasible (the m-array would be of dimensions 13,938 x 13,939, i.e., >194 million cells). We therefore applied only the END model.

Based on the biology of storks, we first made the following assumptions about the demographic processes. Survival differs strongly between juveniles and older individuals (Schaub et al., 2004) and was therefore assumed to be structured into two age classes (juvenile and adult survival). Recruitment into the breeding population was assumed to occur between the age of 1 year (i.e., 2^nd^ calendar year) and the age of 4 years, with all individuals assumed to have recruited by age of 5 years, consistently with the literature (Muller et al., 2026). We assumed survival and resighting probability to vary over time, while recruitment and natal dispersal were assumed to be temporally constant. Survival, recruitment and resighting probability were assumed to vary spatially (among hexagons).

To construct the END-arrays used in our model, we filtered the capture–recapture data to include only resightings of birds observed as breeders, because individuals not observed breeding cannot be reliably classified as non-breeders. In the rare cases where a bird was reported as breeding in more than one spatial unit within the same year, we retained only the first observation. For each hexagon, we summarized the filtered capture–recapture data into END-arrays. The juvenile-END-array contained birds released as juvenile, and the breeder-END-array only birds released as breeder. To avoid bias due to occasional breeding dispersal, breeders whose last observation in hexagon *i* was followed by a later breeder observation in another hexagon were excluded from the “never re-encountered” category of the row of the breeder-END-array, i.e., were censored.

The END model was fitted to the data using the likelihood described above (eqn. 12). To reduce the number of deterministic nodes and repeated operations in the model, several calculations, including the multinomial likelihood contributions, were embedded in custom NIMBLE functions. In addition, different NIMBLE functions were used depending on the release year to avoid unnecessary computations for years with fewer possible future encounter occasions. Rows in END-arrays where no individual was released were removed from the likelihood as they do not contribute any information to inference. For more details, see the code provided.

The hierarchical structure of the parameters was specified as follows. Temporal variability in survival and resighting probability was modeled using a temporal random effect around a hexagon-dependent mean:

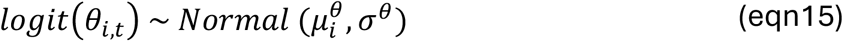

where *θ*_*i,t*_ is the parameter of interest, 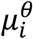 the hexagon-dependent mean of that parameter, and *σ*^*θ*^ the standard deviation of the temporal random effect (Supplement: Table S4 for symbol definitions).

To model spatial variability in mean survival, mean resighting probability and recruitment, we accounted for spatial autocorrelation using an intrinsic autoregressive (ICAR) prior (Besag, 1974), as previously applied to White Stork demographic parameters in Germany (Muller et al., 2026):

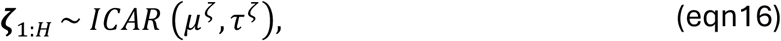

where *μ*^*ζ*^ is the overall mean of the spatially varying parameter *ζ, H* = 101 the number of spatial units, and *τ*^*ζ*^ is the precision of the ICAR distribution, which controls the strength of the spatial autocorrelation. More information about ICAR distributions is available in Appendix S1.

To model natal dispersal, we used a more flexible version of the distance-based approach considered in simulation scenarios e–h. Instead of a simple exponential distance kernel, we used a more flexible power-exponential kernel, which included two parameters: *α*_*d*_, which controls the spatial scale of dispersal, and *β*_*d*_, which controls the shape of the distance-decay relationship. We calculated the distance *d*_*i,j*_ (standardized between 0 and 1) between the centroids of our hexagons; when a hexagon was not entirely within Germany, we took the centroid of its surface inside Germany only. To test for conspecific attraction, which often plays a role in natal dispersal (Buxton et al., 2020), we included the mean population size in the destination hexagon across the study period, *P*_*j*_, as a covariate on the log-scale. *P*_*j*_ was obtained from surveys conducted at the municipality (Kreis) level in Germany; more information is available in Appendix S1 for how Kreis data were aggregated into hexagons. Finally, we included residual noise *η*_*i,j*_ ∼ *Normal* (0, *σ*^*η*^) to account for variation not explained by distance and population size. We had thus:

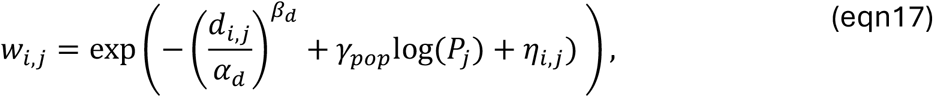

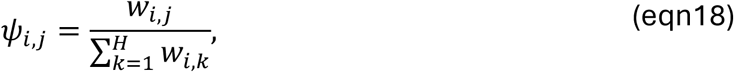

where *γ*_*pop*_ is the effect of destination population size on natal dispersal probability.

We tested for the presence of immediate trap-response, a common feature in capture-mark-recapture analyses, using the test 2.CT from the R package R2Ucare (Gimenez et al., 2018) on our adult breeder resighting data, stratified by hexagon (see Appendix S1). This test found a significant trap-happiness effect, meaning that a breeder resighted at year *t* − 1 was more likely to be resighted at year *t* than a breeder not resighted at year *t* − 1. To account for that effect, we modeled resighting probability of breeders with *p*_*x,i,t*_, *x* denoting whether an individual had been resighted in the previous year (*x* = 1) or not (*x* = 2), and *p*_1, *i,t*_ and *p* _2, *i,t*_ were estimated as separate fixed effects.

We assigned weakly informative priors to all unknown stochastic parameters (Supplement: Table S2). We ran the model in NIMBLE for 40,000 iterations, with a burn-in of 5,000 and a thinning of 20, over 4 chains.

From the estimated recruitment probabilities, we derived the mean age at first breeding in each hexagon (Appendix S1). We used data on the number of juveniles produced in each hexagon to derive absolute number of juvenile dispersers between hexagons. These data were available at the municipality level, as for population size, and were aggregated to the hexagon level (Appendix S1). The vector of the number of juveniles ***J***_*i,t*_ that survived and settled in each destination hexagon *j*, or died, was modelled as

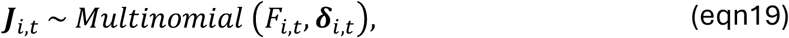

where *F*_*i,t*_ is the number of juveniles produced in hexagon *i* and 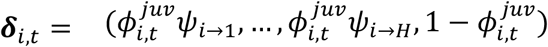. The number of immigrants in hexagon *j* in year *t* (*I*_*j,t*_) was calculated as the sum of juveniles originating from all other hexagons and settling in *j*, whereas the number of emigrants from hexagon *i* in year *t* (*E*_*i,t*_) was calculated as the sum of juveniles originating from *i* and settling in all other hexagons. Net dispersal was defined as the difference between immigrants and emigrants in each hexagon.

We derived source–sink metrics following Runge et al. (2006). For each hexagon *i* and year *t*, we computed the intrinsic growth rate *C*_*i,t*_, using the expected numbers of recruiting immigrants 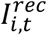 and emigrants 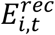 (Appendix S1):

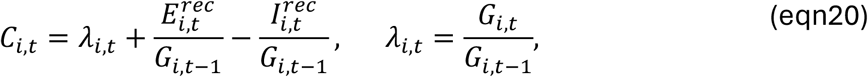

where *G*_*i,t*_ is the number of breeding individuals in hexagon *i* at time *t*.

Because the number of juveniles produced before the study period was unknown, 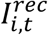 and 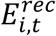 were incomplete during 2001–2004, when individuals born before the start of the study could still have recruited. To obtain the mean intrinsic growth rate per hexagon 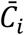, we therefore calculated the geometric mean of *C*_*i,t*_ over 2005–2023, the period for which all recruitment ages considered in the model could be represented. Hexagons with 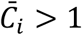 were classified as sources, whereas those with 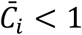 were classified as sinks.

## Results

### Simulation study

Across the first eight simulation scenarios (a-h), bias was small for survival and resighting probability, and similar in the multistate and END models (Fig. 1), as were the corresponding parameter estimates (Supplement: Fig. S4). On average, fidelity showed a negative bias when dispersal was modeled with the Dirichlet distribution, and natal dispersal showed a positive bias (Fig. 1a-d). Biases declined with increasing number of marked individuals but increased with increasing number of spatial units (Fig. 1a-d). Bias was relatively low in all scenarios where dispersal was modeled as a function of distance (Fig. 1e-h). Here, bias (negative for fidelity and positive for dispersal) declined with increasing number of marked individuals, but in contrast to the model with Dirichlet priors it also declined with increasing number of spatial units (Fig. 1e-h). Across the first eight simulation scenarios, RMSE and coverage showed the same overall patterns as bias (Supplement: Figs. S5-S6).

**FIGURE 1.**
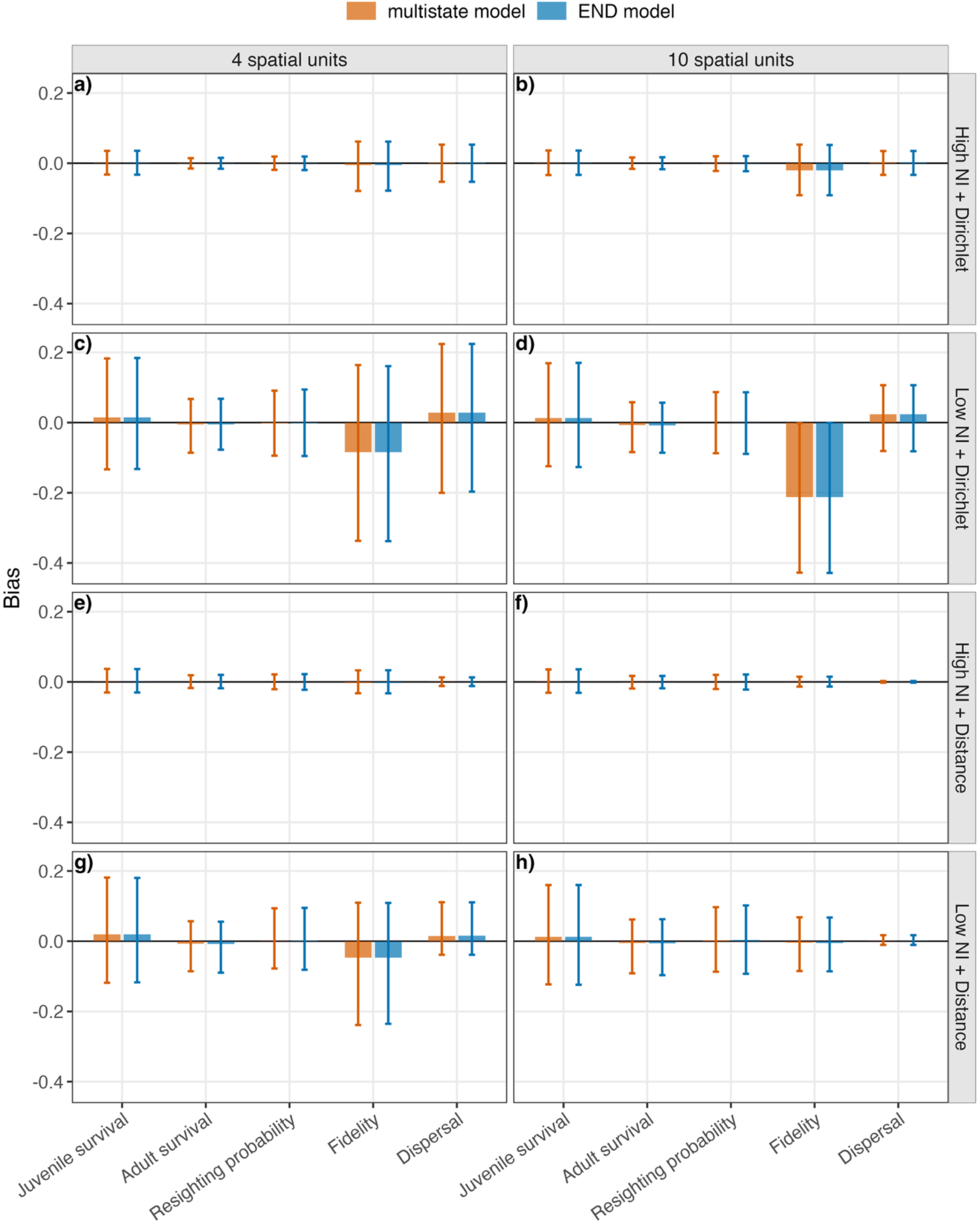
Bias, calculated as the posterior mean minus the true value, in the multistate and END models across eight simulation scenarios (a-h) varying in the number of spatial units (4 or 10), number of released individuals (NI; high [100 per spatial unit, year and age class] or low [5 per spatial unit, year and age class]), and natal dispersal model (Dirichlet or distance). Values were calculated for each parameter element (e.g., survival in spatial unit 1 for survival) and run; plotted values show the mean across all parameter elements and runs within each parameter category, and error bars show the corresponding 2.5th and 97.5th percentiles. “Fidelity” denotes remaining in the natal spatial unit, whereas “dispersal” denotes natal dispersal between different spatial units.

In scenario i, including temporal variation in all parameters, the END model also tracked the multistate model closely when results were examined year by year (Supplement: Figs. S7-S10). The last year showed the known estimation issue in the presence of a temporal fixed effect (Lebreton et al., 1992; Supplement: Figs. S8-S10). In scenario j, including 3 age classes and delayed recruitment, all parameters were estimated similarly and with little bias in both the multistate and END models (Supplement: Fig. S11). However, for recruitment, the RMSE was on average slightly higher in the END model, indicating greater uncertainty (Supplement: Fig. S11c). Finally, in scenarios where dispersal was modeled as a function of distance, the associated hyperparameter *α*_*+*_ was on average biased upwards due to its broad prior, but this bias strongly decreased as the number of marked individuals and the number of spatial units increased (Supplement: Fig. S12).

Computational (MCMC) efficiency differed strongly among models. In all scenarios, the END model was consistently more efficient than the multistate model (Fig. 2). The gain increased with the number of spatial units: efficiency was an order of a magnitude higher in the END model compared to the multistate in the 4 spatial units scenarios but about 20 times higher in the 10 spatial units scenarios. Computational efficiency was also higher for the END model when temporal variability was included (Supplement: Fig. S13). The largest gain among all tested settings was observed in the model with three age classes and delayed recruitment, where the END model was on average more than 30 times more efficient than the multistate model (Supplement: Fig. S14).

**FIGURE 2.**
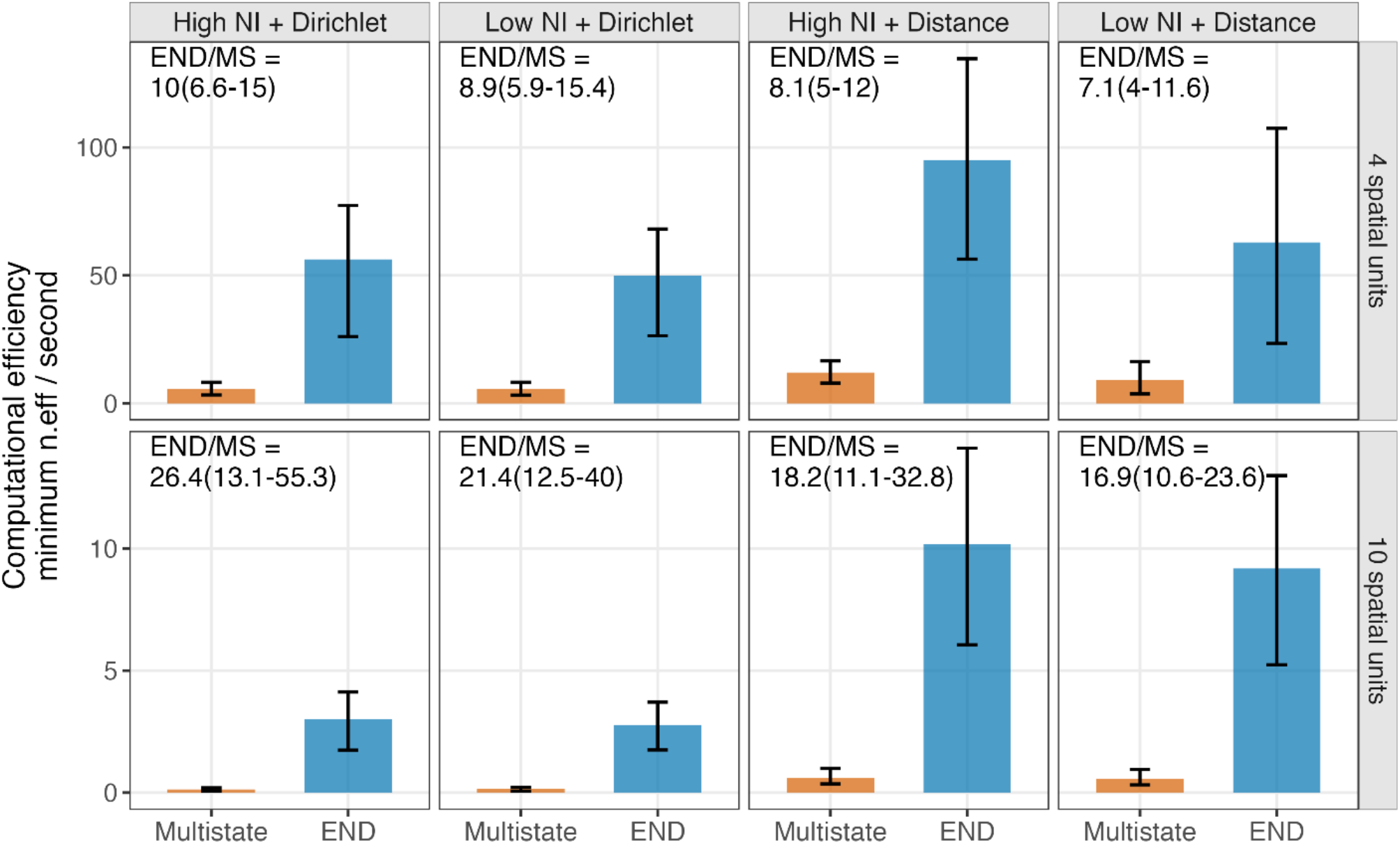
Computational efficiency, calculated as the minimum effective sample size divided by runtime, in the multistate (MS) and END models across eight simulation scenarios varying in the number of spatial units (4 or 10), number of released individuals (NI; high or low), and natal dispersal formulation (Dirichlet or distance). Plotted values show the mean across runs and error bars show the 2.5^th^ and 97.5^th^ percentiles across runs; labels give the mean END/MS efficiency ratio with its 2.5^th^ and 97.5^th^ percentiles across runs.

### Case study

Convergence, assessed using the Gelman–Rubin statistic (Brooks & Gelman, 1998), was good for the quantities of interest (max.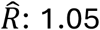; Supplement: Fig. S16a). Effective sample sizes were satisfactory in all quantities of interest, although lower for *α*_*d*_, *β*_*d*_ and *σ*_*η*_ (min. 276; Supplement: Fig. S16b).

We found marked spatial variation among the 101 hexagonal spatial units in Germany in survival and recruitment. Juvenile survival was generally highest in hexagons located in the west of the country, whereas lower values were observed in the north-east (Fig. 3a). Adult survival also showed a clear spatial pattern, being higher in the south and south-west and exhibiting lower values towards the north-east (Fig. 3b). Age at first breeding occurred earlier in western and south-western hexagons than in north-eastern parts of the study area (Fig. 3c).

**FIGURE 3.**
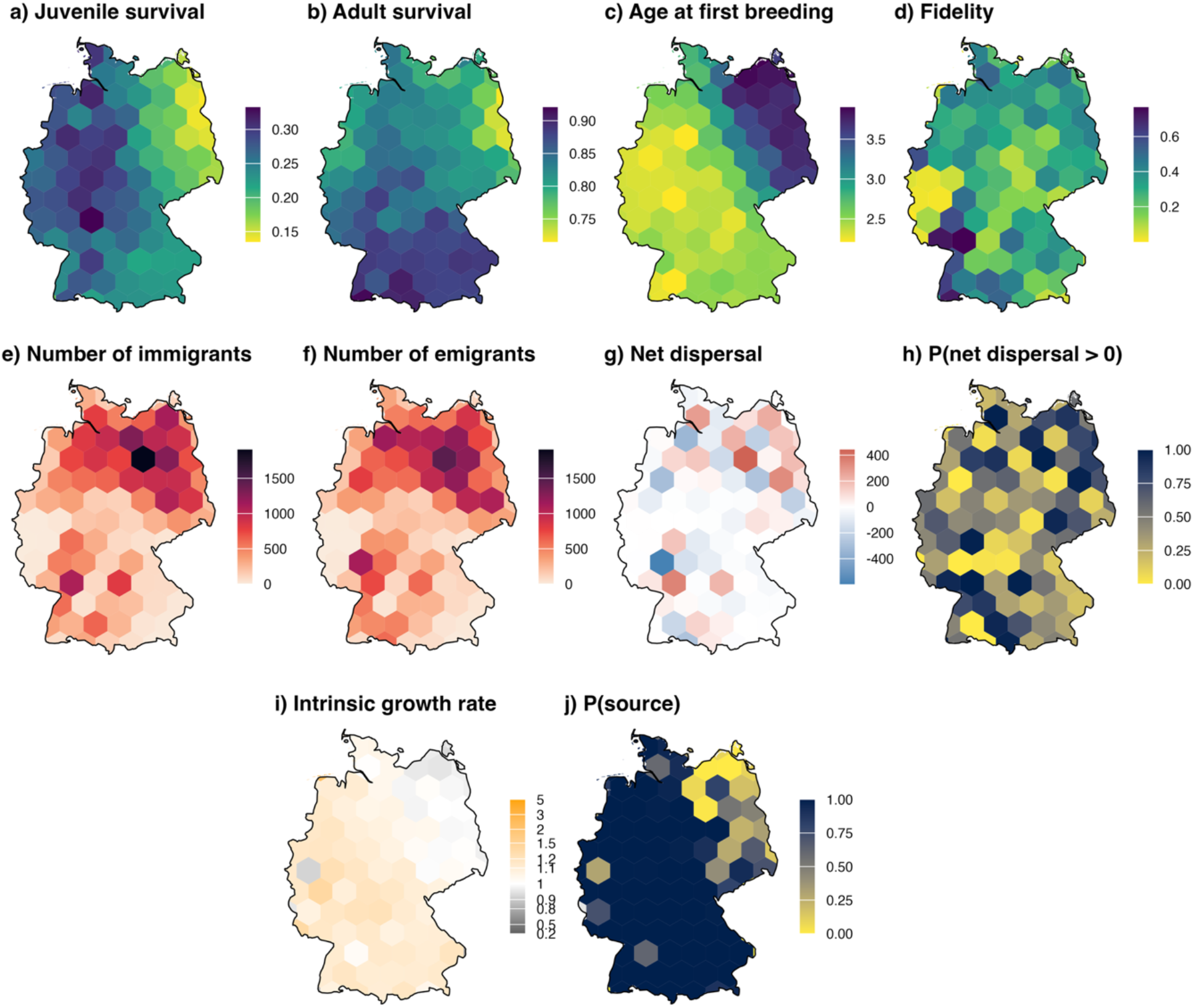
Spatial variation among the 101 hexagonal spatial units covering Germany in posterior means and posterior probabilities of key demographic and dispersal quantities obtained from the model. Maps show, from left to right and top to bottom: a) posterior mean juvenile survival, b) posterior mean adult survival, c) posterior mean age at first breeding, d) posterior mean fidelity, e) posterior mean number of immigrants, f) posterior mean number of emigrants, g) posterior mean net dispersal (immigrants minus emigrants), h) posterior probability that net migration is positive, i) posterior mean intrinsic growth rate from 2005 to 2023, and j) posterior probability of being a source.

Dispersal probability declined strongly with distance (Supplement: Fig. S17), with *α*_*+*_ = 0.027 (95% CrI: 0.022–0.032). *β*_*d*_ = 0.660 (95% CrI: 0.623–0.700) indicated a fat-tailed decline, consistent with predominantly short-distance dispersal but with some longer-distance movements. The estimated median dispersal distance was 103.81 km (95% CrI: 98.79–109.19), while 5% of individuals dispersed more than 374.73 km (95% CrI: 352.70-399.06). The effect of population size at hexagon of destination was positive, with *γ*_BCB_ = 0.775 (95% CrI: 0.711–0.838), indicating that hexagons with larger mean population sizes had higher probabilities of receiving natal dispersers. Residual variation among origin– destination hexagon pairs remained substantial, with *σ*_*η*_ = 0.661 (95% CrI: 0.605–0.722), showing that dispersal probabilities still varied beyond the effects of distance and destination population size (see Supplement: Fig. S18 for per-hexagon dispersal probabilities).

Fidelity showed a heterogeneous spatial pattern, with no clear large-scale gradient across Germany (Fig. 3d). The posterior mean numbers of immigrants and emigrants per hexagon were often similar (Fig. 3e-f). The posterior net dispersal was therefore often close to zero, but in north-eastern Germany, net dispersal tended to be positive in most hexagons, while we found a more heterogeneous pattern in the south-western part of our study area (Fig. 3g-h). Finally, most spatial units in Germany were estimated to act as sources. The majority of the estimated sinks were located in northeastern Germany (Fig. 3i-j).

## Discussion

We developed a novel multistate capture-recapture model to estimate natal dispersal among discrete spatial units from capture-recapture data, while jointly accounting for spatial variation in demographic parameters and imperfect detection, but at a much lower computational cost than conventional multistate capture–recapture models. Our simulation study revealed that our novel model yielded parameter estimates very similar to those of a conventional multistate model, while greatly improving computational efficiency, especially when the number of spatial units is high. This opens the way for analyzing spatially structured capture–recapture data in systems where conventional multistate models would be computationally impossible to apply, and for reconstructing spatially detailed patterns of natal dispersal across large landscapes, as illustrated by our case study modelling dispersal across 101 spatial units in Germany.

In recent years, several approaches have been proposed to improve the computational efficiency of multistate models, including marginalization (Yackulic et al., 2020), optimization of the multinomial likelihood (Muller & Badia-Boher, 2026), and the use of Hamiltonian Monte Carlo to improve MCMC sampling efficiency (Hollanders, 2026). However, these approaches alone remain insufficient for modelling natal dispersal because they do not incorporate the two key simplifications underlying the efficiency of the END model: the separate treatment of juveniles and breeders (the latter assumed to remain in the same spatial unit) and, within the juvenile component, the aggregation of resightings across years for individuals re-encountered outside their natal spatial unit. Therefore, the END model represents the first capture-recapture framework specifically designed to address the computational challenges of estimating natal dispersal across many spatial units.

Our simulations showed that the END model did not result in estimates of survival and natal dispersal with increased bias or reduced precision in, despite aggregating resightings of dispersing individuals across all subsequent occasions. This is possible because natal dispersal occurs only once, during the first year after marking. A small loss of precision was observed only for the recruitment probability. This was expected because recruitment cannot be fully estimated without information on the timing of resightings and is therefore informed primarily by individuals that remain in their natal spatial unit.

Another insight from our simulations is that dispersal estimates were sensitive to the prior used for dispersal probabilities. With weakly informative priors, fidelity tended to be underestimated and dispersal probabilities (excluding fidelity) overestimated, especially when the number of marked individuals was low. However, the effect of the number of spatial units differed between dispersal formulations. In Dirichlet-based scenarios, bias increased with the number of spatial units because the prior assigns the same expected probability to all destinations, thereby placing decreasing prior weight on fidelity as the number of units increases (Kotz et al., 2000). This indicates that these posteriors are sensitive to the prior. In contrast, in distance-based scenarios, bias decreased with the number of spatial units, likely because additional units improved estimation of the dispersal kernel; bias was then small even when the number of marked individuals was low. These results suggest that weakly informative Dirichlet priors should be avoided when the number of spatial units is large (i.e., ≥10). In such cases, structured dispersal models based on distance and/or spatial covariates are preferable, because they can improve estimation while providing biologically interpretable information on the drivers of natal dispersal.

We applied our new model to white stork capture–recapture data across Germany, divided into 101 hexagons. We found marked spatial variation in demographic parameters, with both juvenile and adult survival being lower in the north-eastern part of Germany than in the south-western part of the country, and age at first breeding occurring earlier in the south-west. This pattern is consistent with recent findings (Muller et al., 2026). The estimated dispersal kernel (Supplement: Fig. S17) followed a power-exponential form with a heavier tail than a simple exponential, suggesting that most individuals disperse over short distances, whereas a smaller proportion disperse much farther, as reported in many bird species (Fandos et al., 2023). The median dispersal distance estimated here exceeded previous estimates for white storks (Chernetsov et al., 2006; Ječmenica & Kralj, 2017), likely because our larger study area captured more long-distance movements. In addition, population size in the destination spatial unit had a strong positive effect on immigration probability, suggesting conspecific attraction in natal dispersal (Buxton et al., 2020), a pattern not previously documented in white storks, to our knowledge. Going one step further, we translated dispersal probabilities into juvenile fluxes using productivity data. This made it possible to estimate net dispersal and intrinsic growth rate – used to assess whether a given unit acted as a source or a sink, a challenge at such a large scale (Furrer & Pasinelli, 2016). In this application, some sharp contrasts between neighbouring hexagons may reflect uneven ringing effort within hexagons and the proportional allocation of Kreis-level population sizes among hexagons. Nevertheless, immigration and emigration were overall balanced and most spatial units acted as a source, whereas some spatial units in the north-eastern part of Germany appeared to receive more immigrants than they produced emigrants and to act as a sink, in line with the less favorable demographic parameters estimated in this region. Overall, this case study illustrates the important potential of our approach to improve understanding of spatial population dynamics over large geographic scales, and also to provide conservation-relevant information by identifying areas with unfavourable demographic rates that may require targeted management.

Our framework could be extended in several directions. First, dispersal out of the study area could be represented by extending the dispersal kernel beyond the study boundaries (Badia-Boher et al., 2023); accounting for this emigration would prevent potential downward bias in juvenile survival estimates near the boundaries (Schaub & Royle, 2014). Second, breeding dispersal could be incorporated. Although negligible in many species and therefore often safely ignored, it can bias natal dispersal estimates when substantial, because dispersal inferred from the natal location to a subsequent breeding location may reflect natal dispersal, breeding dispersal, or both. Extending our framework to incorporate breeding dispersal would, however, require assumptions restricting possible destinations, for example to neighbouring spatial units or units within a maximum distance, to avoid recovering the dense structure of a conventional multistate model. Finally, integrating dead-recovery data into our framework could be another extension. It would provide information on true survival and help distinguish mortality from permanent emigration from a given spatial unit (Burnham, 1993).

In conclusion, we developed a computationally efficient capture–recapture model for reconstructing natal dispersal networks across many spatial units while jointly accounting for spatial variation in demographic parameters. Although our case study used hexagonal spatial units, the framework is equally applicable to capture-recapture data with other discrete spatial structures, such as breeding colonies, management units or large metapopulation systems. Further work should integrate our framework with other modeling approaches, for example through an integrated population model (IPM; Schaub & Kéry, 2022), to provide a complete picture of population dynamics across space. More broadly, such dispersal-network-based approaches could help quantify the spatial extent of demographic influence, that is, how much local demographic processes such as survival, dispersal and productivity affect population dynamics at other locations. This would be particularly valuable for conservation, by helping move from local management actions to spatially coherent strategies that account for how demographic processes are connected across populations.

## Acknowledgments

We thank Marc Kéry and Jaume A. Badia-Boher for providing helpful advice and comments. This work was supported by the Swiss National Science Foundation through grant no. 215689 awarded to Michael Schaub.

## Author contributions

Matia H. Muller and Michael Schaub conceived the ideas and designed methodology; Christof Herrmann, Wolfgang Fiedler and Olaf Geiter collected and stored the data used in the case study; Matia H. Muller, with the help of Fabian R. Ketwaroo and Michael Schaub, analysed the data; Matia H. Muller led the writing of the manuscript. All authors critically revised earlier drafts and gave final approval for publication.

## Conflict of interest

We declare no conflict of interest.

## Data availability

Upon acceptance, data and code will be archived in a permanent public repository. In the meantime, data can be requested by contacting the corresponding author by email.

## Notes

### Competing Interest Statement

The authors have declared no competing interest.

