## Supplement for "Efficient capture–recapture inference for spatially varying natal dispersal, survival and recruitment"

**SUPPLEMENTARY FIGURES AND TABLES**

In this document additional figures and tables are provided:

- Figures S1 and S2 provide supplementary information about the aggregation of multistate capture-histories in conventional m-arrays and END-arrays.
- Figures S3 to S14 provide supplementary information about the simulation study and its results.
- Figures S15 to S18 provide supplementary information about the case study and its results.
- Table S1 provides details about notation used in the formulation of the conventional multistate and END models.
- Table S2 provides the predefined distributions used to simulate capture-histories under the different scenarios.
- Table S3 provides details about notation used in the simulation study.
- Table S4 provides details about notation used in the case study.
- Table S5 provides the priors assigned to stochastic parameters in the White stork case study.

Capture-histories:

Y1 Y2 Y3

ID1: 1 4 4
ID2: 2 6 6
ID3: 1 4 0
ID4: 5 0 5
ID5: 3 0 0
ID6: 0 3 6

States:

1 = juvenile in unit A
2 = juvenile in unit B
3 = juvenile in unit C
4 = adult in unit A
5 = adult in unit B
6 = adult in unit C

M-array (multistate)

| | Y2 S1 | Y2 S2 | Y2 S3 | Y2 S4 | Y2 S5 | Y2 S6 | Y3 S1 | Y3 S2 | Y3 S3 | Y3 S4 | Y3 S5 | Y3 S6 | ∅ |
|---|---|---|---|---|---|---|---|---|---|---|---|---|---|
| Y1S1 | 0 | 0 | 0 | 1+1 | 0 | 0 | 0 | 0 | 0 | 0 | 0 | 0 | 0 |
| Y1S2 | 0 | 0 | 0 | 0 | 0 | 1 | 0 | 0 | 0 | 0 | 0 | 0 | 0 |
| Y1S3 | 0 | 0 | 0 | 0 | 0 | 0 | 0 | 0 | 0 | 0 | 0 | 0 | 1 |
| Y1S4 | 0 | 0 | 0 | 0 | 0 | 0 | 0 | 0 | 0 | 0 | 0 | 0 | 0 |
| Y1S5 | 0 | 0 | 0 | 0 | 0 | 0 | 0 | 0 | 0 | 0 | 1 | 0 | 0 |
| Y1S6 | 0 | 0 | 0 | 0 | 0 | 0 | 0 | 0 | 0 | 0 | 0 | 0 | 0 |
| Y2S1 | 0 | 0 | 0 | 0 | 0 | 0 | 0 | 0 | 0 | 0 | 0 | 0 | 0 |
| Y2S2 | 0 | 0 | 0 | 0 | 0 | 0 | 0 | 0 | 0 | 0 | 0 | 0 | 0 |
| Y2S3 | 0 | 0 | 0 | 0 | 0 | 0 | 0 | 0 | 0 | 0 | 0 | 1 | 0 |
| Y2S4 | 0 | 0 | 0 | 0 | 0 | 0 | 0 | 0 | 0 | 1 | 0 | 0 | 1 |
| Y2S5 | 0 | 0 | 0 | 0 | 0 | 0 | 0 | 0 | 0 | 0 | 0 | 0 | 0 |
| Y2S6 | 0 | 0 | 0 | 0 | 0 | 0 | 0 | 0 | 0 | 0 | 0 | 1 | 0 |

**FIGURE S1.** Aggregation of multistate individual capture-histories into a multistate m-array. Y = year, S = state, ∅ = never re-encountered. The capture histories of each individual has a different color and the same colors are used in the m-array table to show where the capture-history data enter the m-array.

Capture-histories:

Y1 Y2 Y3

ID1: 1 4 4

ID2: 2 6 6

ID3: 1 4 0

ID4: 5 0 5

ID5: 3 0 0

ID6: 0 3 6

States:

1 = juvenile in unit A
2 = juvenile in unit B
3 = juvenile in unit C
4 = adult in unit A
5 = adult in unit B
6 = adult in unit C

Juvenile-END-array

| | Y2N | Y3N | UA | UB | UC | ∅ |
|---|---|---|---|---|---|---|
| Y1UA | 1+1 | 0 | 0 | 0 | 0 | 0 |
| Y1UB | 0 | 0 | 0 | 0 | 1 | 0 |
| Y1UC | 0 | 0 | 0 | 0 | 0 | 1 |
| Y2UA | 0 | 0 | 0 | 0 | 0 | 0 |
| Y2UB | 0 | 0 | 0 | 0 | 0 | 0 |
| Y2UC | 0 | 1 | 0 | 0 | 0 | 0 |

Breeder-END-array

| | Y2 | Y3 | ∅ |
|---|---|---|---|
| Y1UA | 0 | 0 | 0 |
| Y1UB | 0 | 1 | 0 |
| Y1UC | 0 | 0 | 0 |
| Y2UA | 0 | 1 | 1 |
| Y2UB | 0 | 0 | 0 |
| Y2UC | 0 | 1 | 0 |

**FIGURE S2.** Aggregation of multistate individual capture-histories into juvenile- and breeder-END-arrays. Y = year, U = spatial unit, N = natal, ∅ = never re-encountered. The capture histories of each individual has a different color and the same colors are used in the m-array table to show where the capture-history data enter the END-arrays.

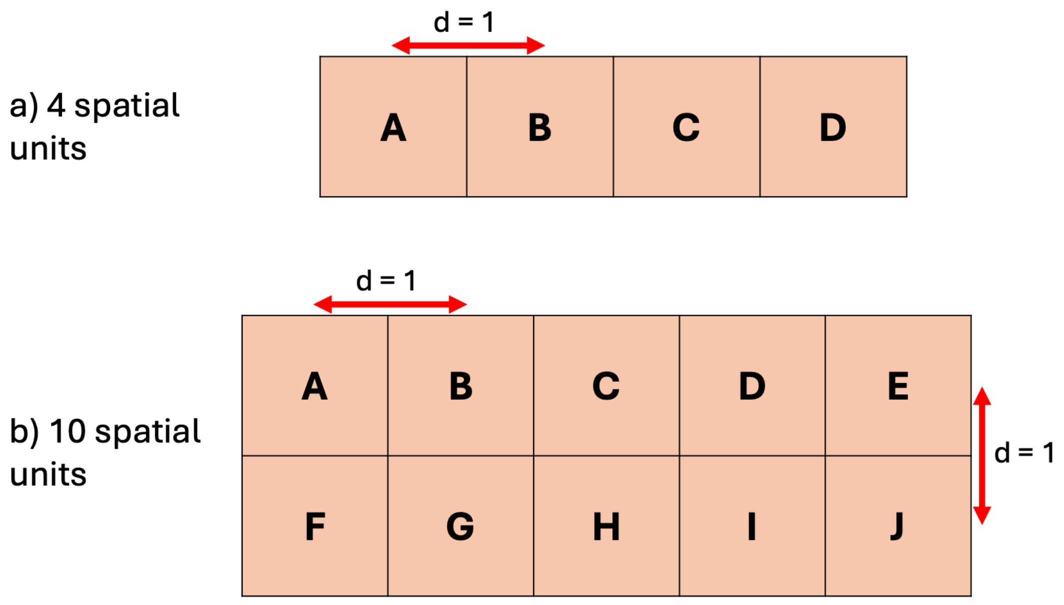


**FIGURE S3.** Spatial arrangement of the spatial units in the simulation study when there are a) 4 spatial units and b) 10 spatial units. The distance (d) between the centers of two adjacent spatial units is always set to 1.

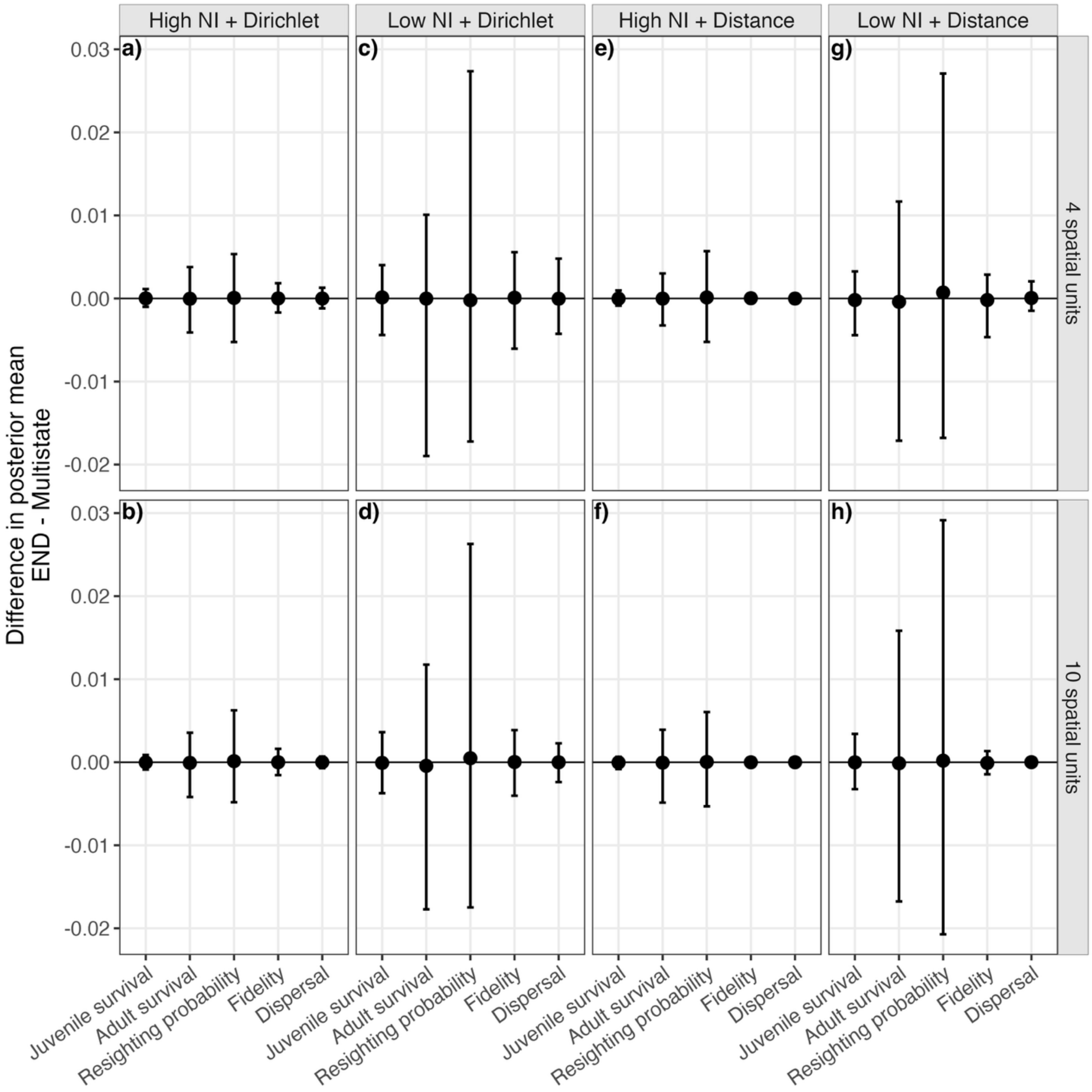


**FIGURE S4.** Difference in posterior mean between the multistate and END models across eight simulation scenarios (a-h) varying in the number of spatial units (4 or 10), number of released individuals (NI; high [100 per spatial unit, year and age class] or low [5 per spatial unit, year and age class]), and natal dispersal model (Dirichlet or distance). Values were calculated for each parameter element (e.g., juvenile survival in spatial unit 1) and run; plotted values show the mean across all parameter elements and runs within each parameter category, and error bars show the corresponding 2.5th and 97.5th percentiles. "Fidelity" denotes remaining in the natal spatial unit, whereas "dispersal" denotes natal dispersal between different spatial units.

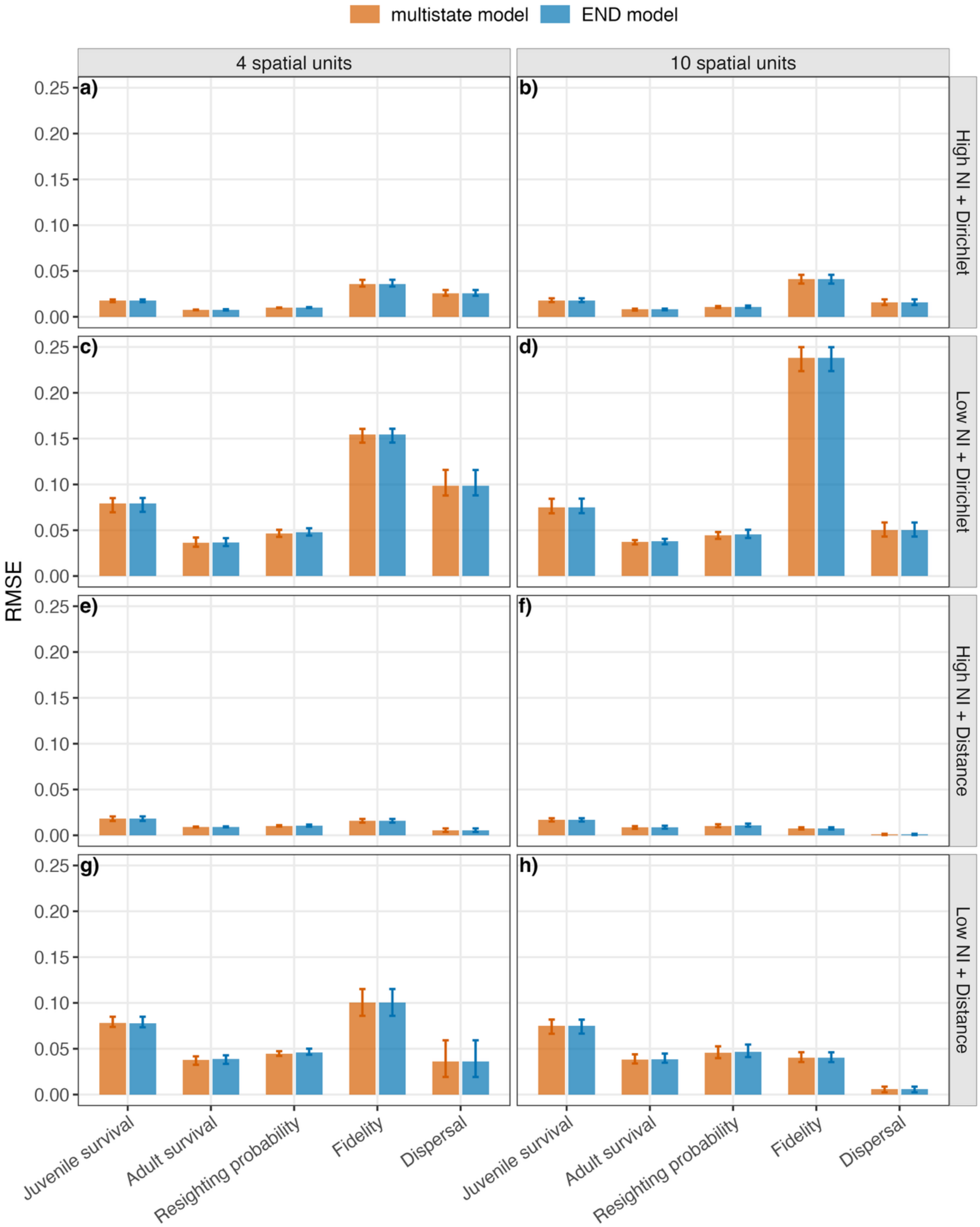

multistate model
END model
4 spatial units
10 spatial units
High NI + Dirichlet
Low NI + Dirichlet
High NI + Distance
Low NI + Distance
RMSE
0.25
0.20
0.15
0.10
0.05
0.00
a)
b)
c)
d)
e)
f)
g)
h)
Juvenile survival
Adult survival
Resighting probability
Fidelity
Dispersal

**FIGURE S5.** Root mean squared error (RMSE) across eight simulation scenarios (a-h) varying in the number of released individuals (NI; high [100 per spatial unit, year and age class] or low [5 per spatial unit, year and age class]), number of spatial units (4 or 10), and natal dispersal model (Dirichlet or distance), and across the two tested models (multistate and END). For each parameter element (e.g., survival in spatial unit 1 for survival), RMSE was calculated across all simulation replicates; displayed values show the mean across parameter elements, and error bars show the 2.5th and 97.5th percentiles across parameter elements. “Fidelity” denotes remaining in the natal spatial unit, whereas “dispersal” denotes natal dispersal between different spatial units.

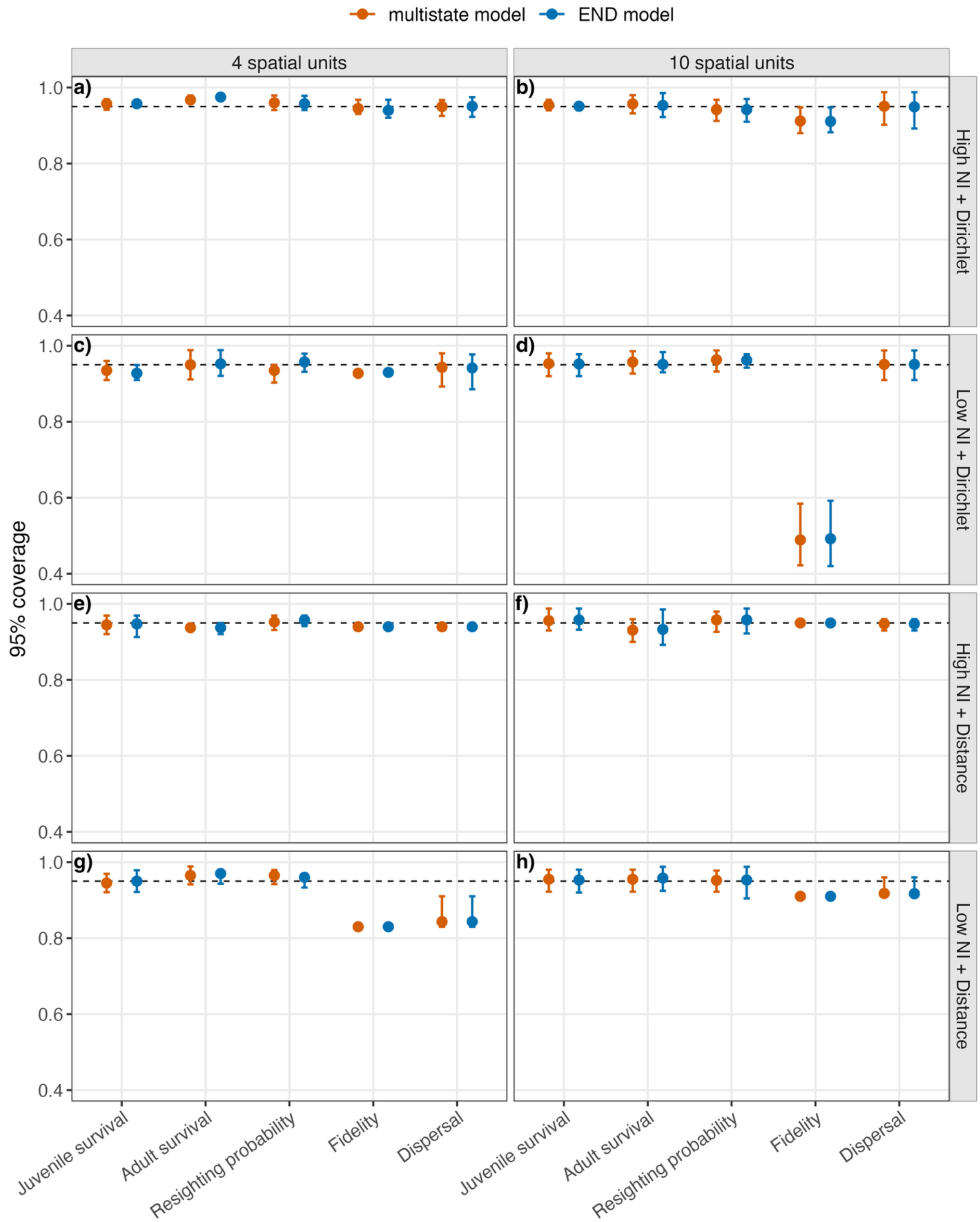


**FIGURE S6.** 95% coverage across eight simulation scenarios (a-h) varying in the number of spatial units (4 or 10), number of released individuals (NI; high [100 per spatial unit, year and age class] or low [5 per spatial unit, year and age class]), and natal dispersal model (Dirichlet or distance), and across the two tested models (multistate and END). For each parameter element (e.g., survival in spatial unit 1 for survival), coverage was calculated across all simulation replicates; points show the mean across parameter elements, and error bars show the 2.5th and 97.5th percentiles across parameter elements. “Fidelity” denotes remaining in the natal spatial unit, whereas “dispersal” denotes natal dispersal between different spatial units. The dashed line shows the expected value of 0.95.

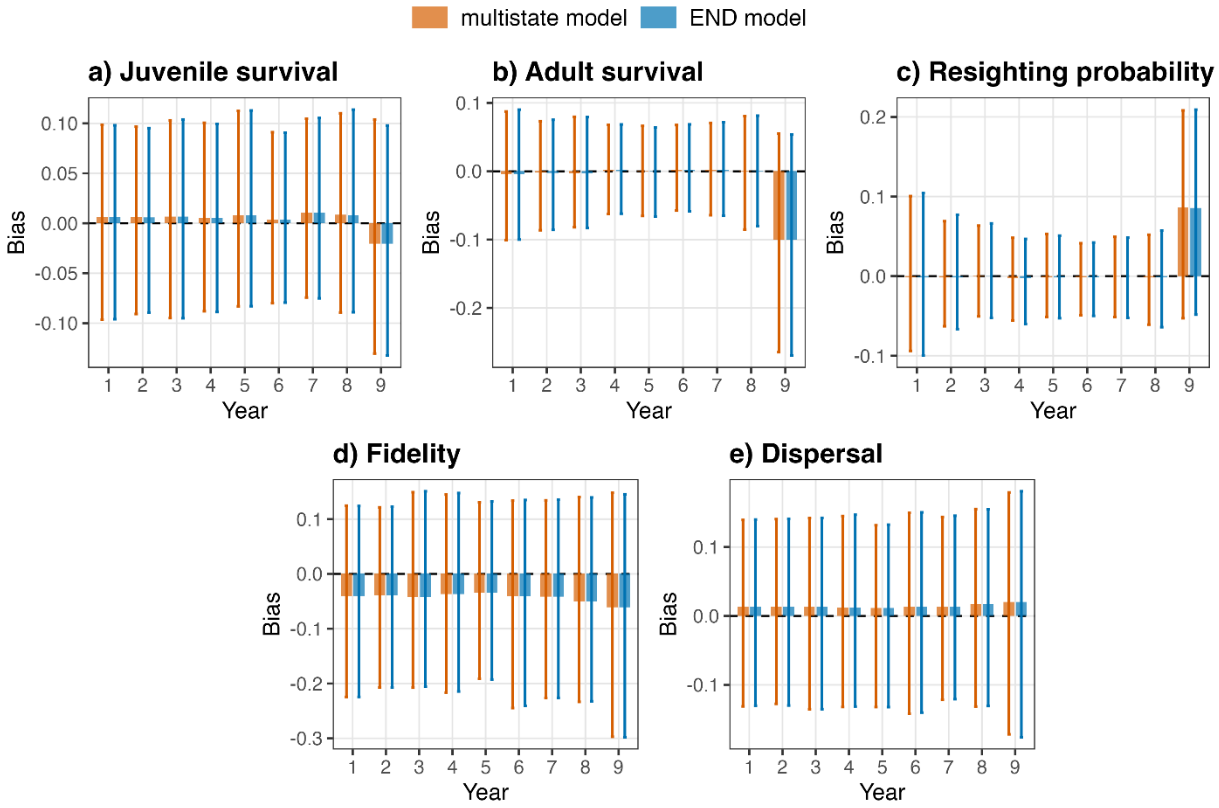


**FIGURE S7.** Per-year bias, calculated as the posterior mean minus the true value, in both END and multistate models under a simulation scenario with temporal variability (scenario i). Values were calculated for each parameter element and run; plotted values show the mean across all parameter elements and runs within each parameter category, and error bars show the corresponding 2.5th and 97.5th percentiles. “Fidelity” denotes remaining in the natal spatial unit, whereas “dispersal” denotes natal dispersal between different spatial units.

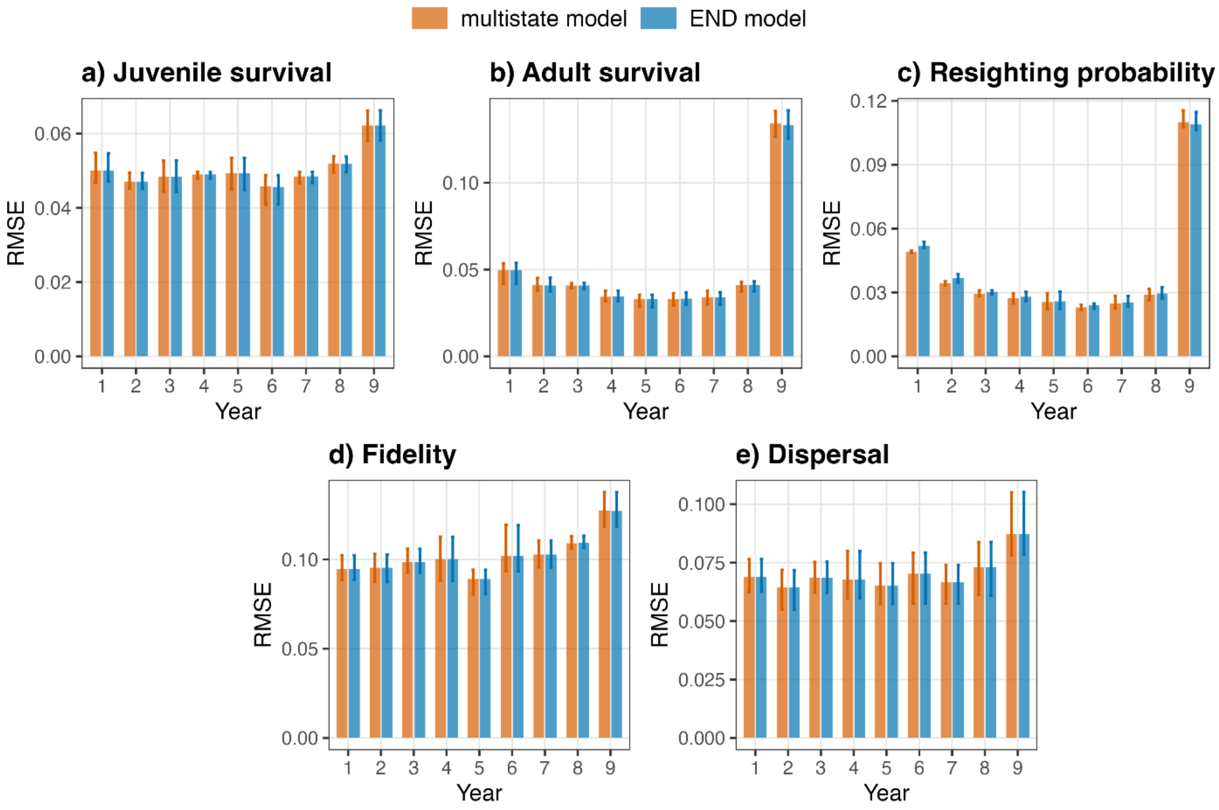


**FIGURE S8.** Per-year root mean squared error (RMSE) in both END and multistate models under a simulation scenario with temporal variability (scenario i). Values were calculated for each parameter element across runs; plotted values show the mean within each parameter category, and error bars show the corresponding 2.5$^{th}$ and 97.5$^{th}$ percentiles. “Fidelity” denotes remaining in the natal spatial unit, whereas “dispersal” denotes natal dispersal between different spatial units.

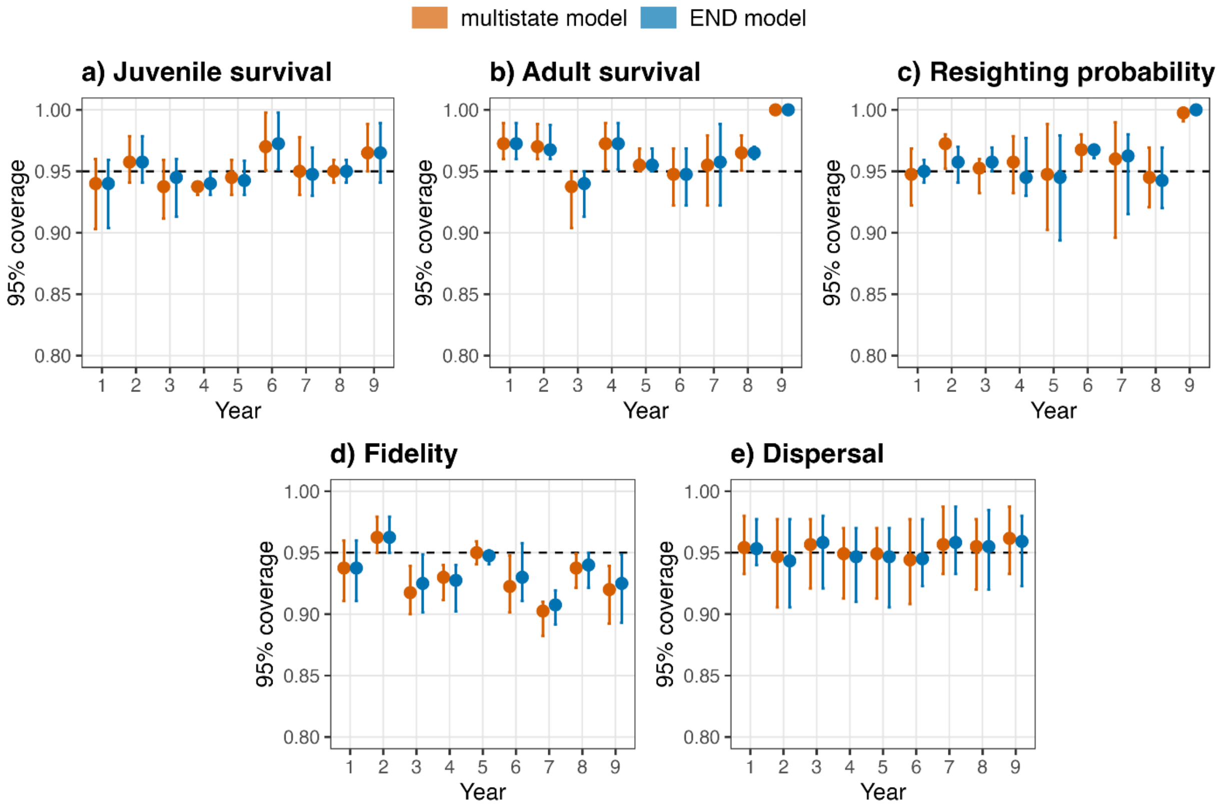


**FIGURE S9.** Per-year 95% coverage in both END and multistate models under a simulation scenario with temporal variability (scenario i). Values were calculated for each parameter element across runs; points show the mean within each parameter category, and error bars show the corresponding 2.5th and 97.5th percentiles. "Fidelity" denotes remaining in the natal spatial unit, whereas "dispersal" denotes natal dispersal between different spatial units.

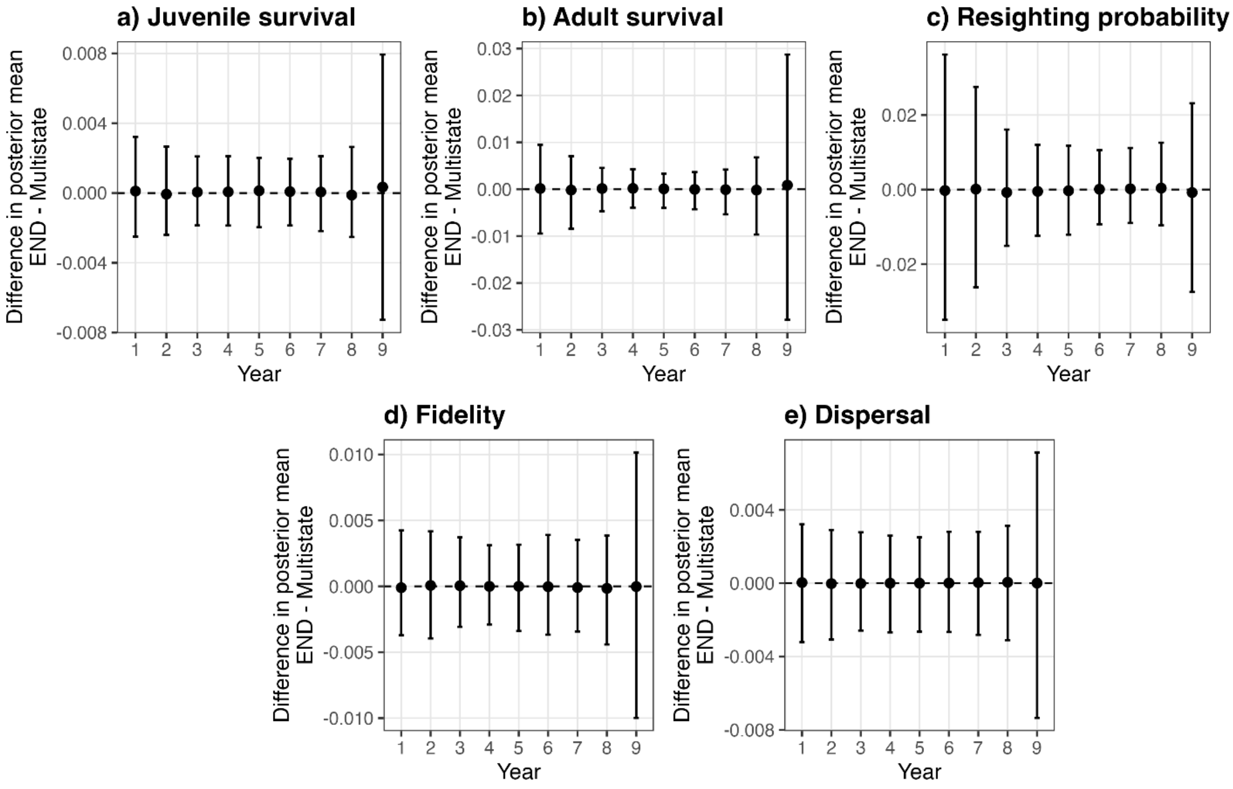


**FIGURE S10.** Per-year difference in posterior means between END and multistate models under a simulation scenario with temporal variability (scenario i). Values were calculated for each parameter element and run; plotted values show the mean across all parameter elements and runs within each parameter category, and error bars show the corresponding 2.5th and 97.5th percentiles. “Fidelity” denotes remaining in the natal spatial unit, whereas “dispersal” denotes natal dispersal between different spatial units.

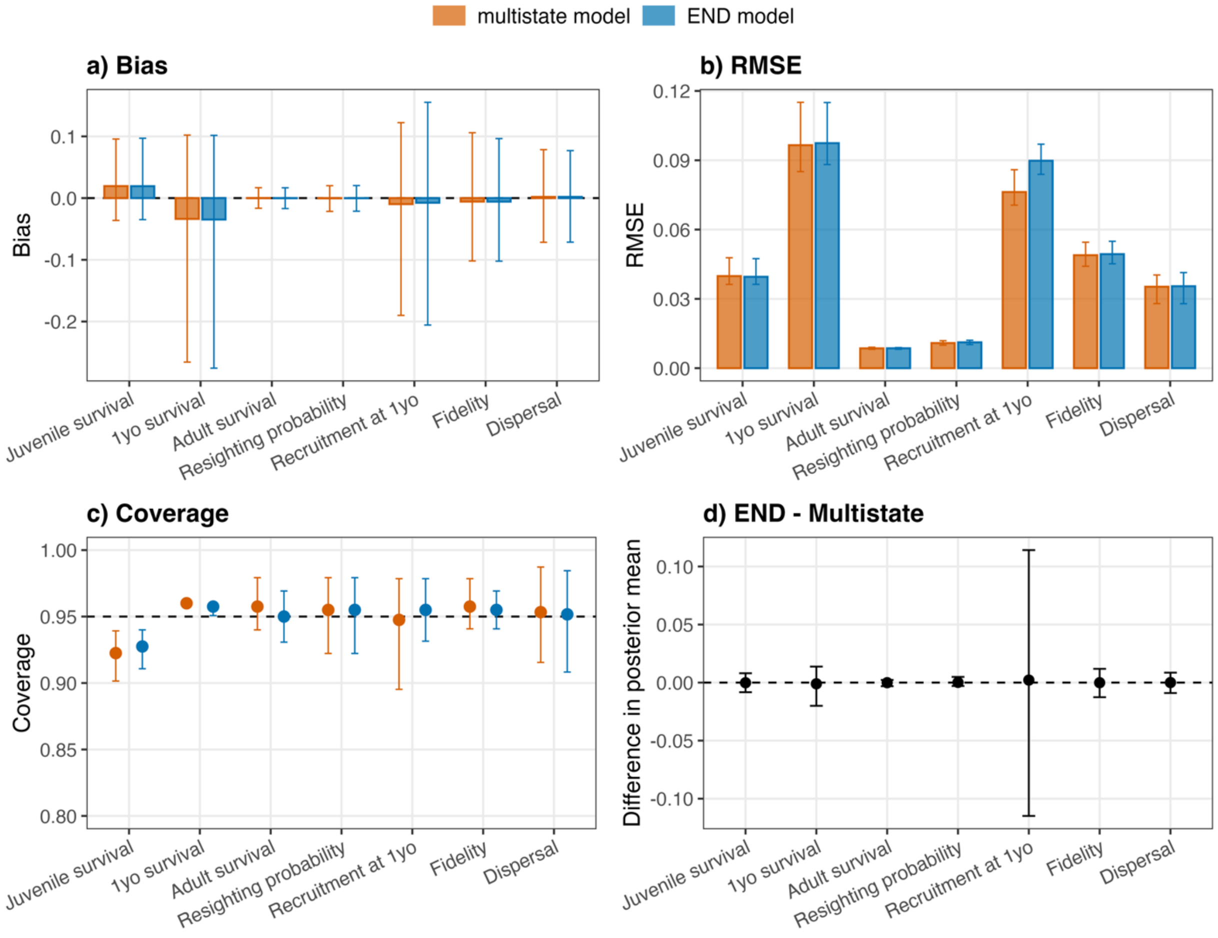


**FIGURE S11.** Performance comparison between the multistate and END models under a simulation scenario with three age classes and delayed recruitment (scenario j). a) Bias, calculated as the posterior mean minus the true value. b) Root mean squared error. c) 95% credible interval coverage. d) Difference in posterior means between END and multistate models applied to the same dataset. In panels a) and d), values were calculated for each parameter element and run; plotted values show the mean across all parameter elements and runs within each parameter category, and error bars show the corresponding 2.5th and 97.5th percentiles. In panels b) and c), values were calculated for each parameter element across runs; plotted values show the mean within each parameter category, and error bars show the corresponding 2.5$^{th}$ and 97.5$^{th}$ percentiles. “Fidelity” denotes remaining in the natal spatial unit, whereas “dispersal” denotes natal dispersal between different spatial units.

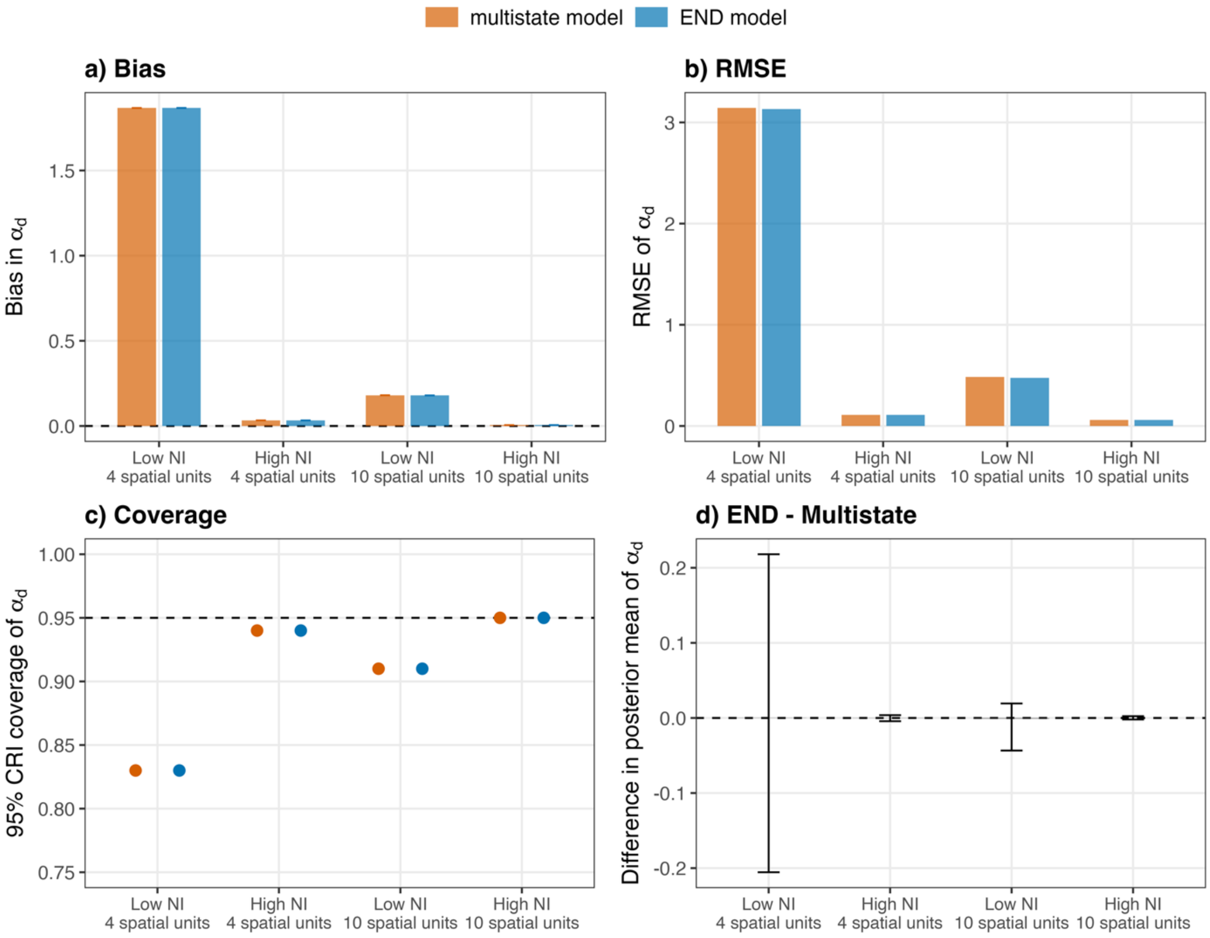


**FIGURE S12.** Comparison between the multistate and END models for the estimation of the parameter $\alpha_d$ under four simulation scenarios varying in the number of marked individuals (NI; high or low) and the number of spatial units (4 or 10). a) Bias in $\alpha_d$, calculated as the posterior mean minus the true value. b) Root mean squared error of $\alpha_d$. c) 95% credible interval coverage of $\alpha_d$. d) Difference in posterior mean of $\alpha_d$ between END and multistate models applied to the same dataset. In panels a) and d), values were calculated for each run; plotted values show the mean and error bars the 2.5$^{th}$ and 97.5$^{th}$ percentiles. In panels b) and c), values are calculated across all runs for each scenario and model.

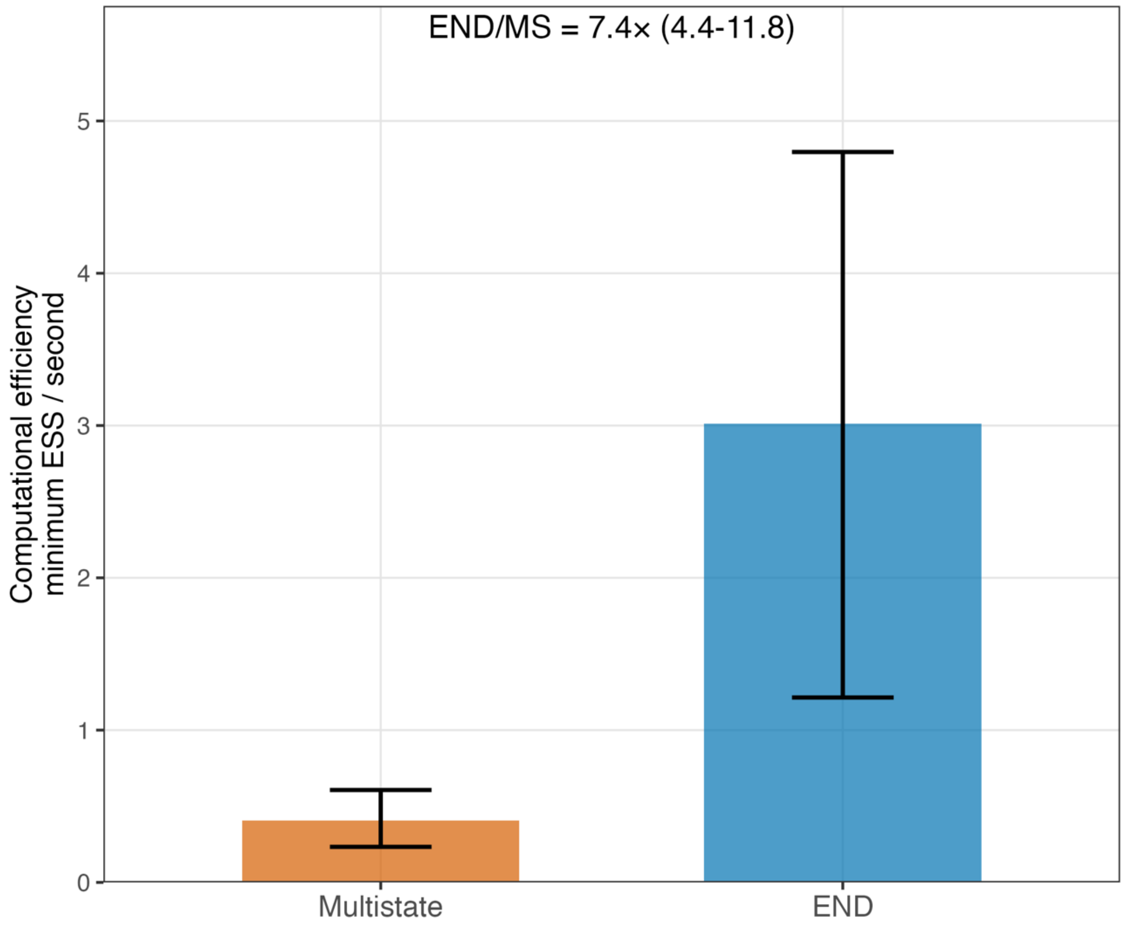


**FIGURE S13**. Computational efficiency, calculated as the minimum effective sample size divided by runtime, in the multistate (MS) and END models under a simulation scenario with temporal variability (scenario i). The last year was excluded because of known convergence issues. Plotted values show the mean across runs and error bars show the 2.5$^{th}$ and 97.5$^{th}$ percentiles across runs; labels give the mean END/MS efficiency ratio with its 2.5$^{th}$ and 97.5$^{th}$ percentiles across runs.

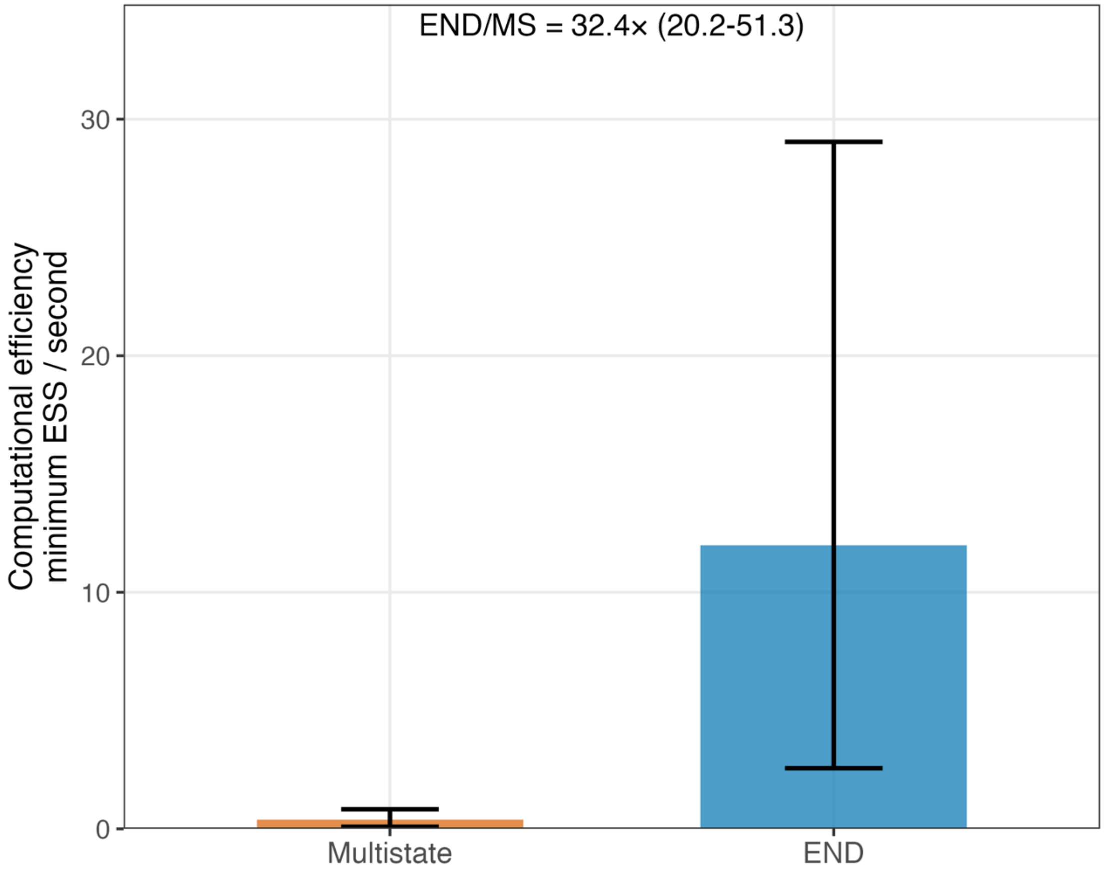


**FIGURE S14**. Computational efficiency, calculated as the minimum effective sample size divided by runtime, in the multistate (MS) and END models under a simulation scenario with three age classes and delayed recruitment (scenario j). Plotted values show the mean across runs and error bars show the 2.5$^{th}$ and 97.5$^{th}$ percentiles across runs; labels give the mean END/MS efficiency ratio with its 2.5$^{th}$ and 97.5$^{th}$ percentiles across runs.

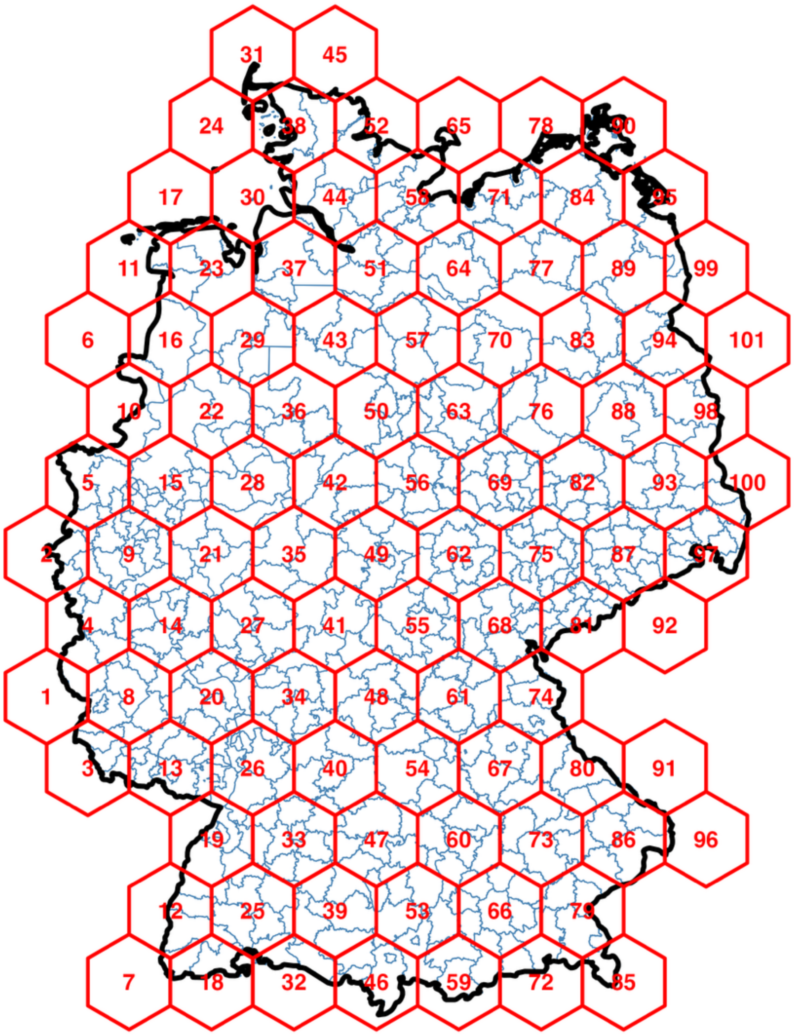


**FIGURE S15.** Spatial discretization of Germany into 101 hexagons used in the case study. Blue lines show German administrative districts ("Kreise"), which were used to compile population size and juvenile data. Red lines show the 101 hexagonal spatial units used in the model, with red numbers indicating the hexagon identifiers.

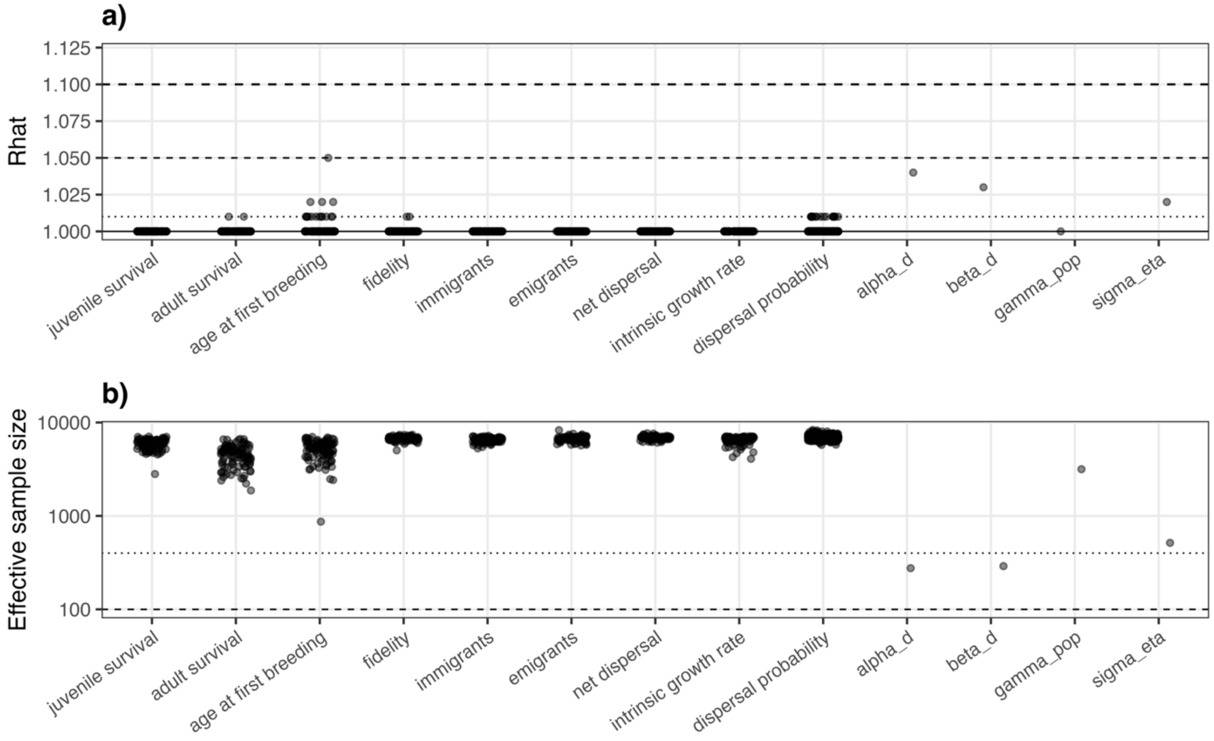


**FIGURE S16.** Convergence diagnostics for parameters and derived quantities of interest in the case study. "Dispersal probability" denotes dispersal to another spatial unit. For each natal spatial unit, diagnostics for dispersal probabilities were restricted to the ten destination units with the highest posterior mean probabilities, thereby excluding the large number of dispersal movements probabilities close to zero. Shown are a) R-hat values and b) effective sample sizes.

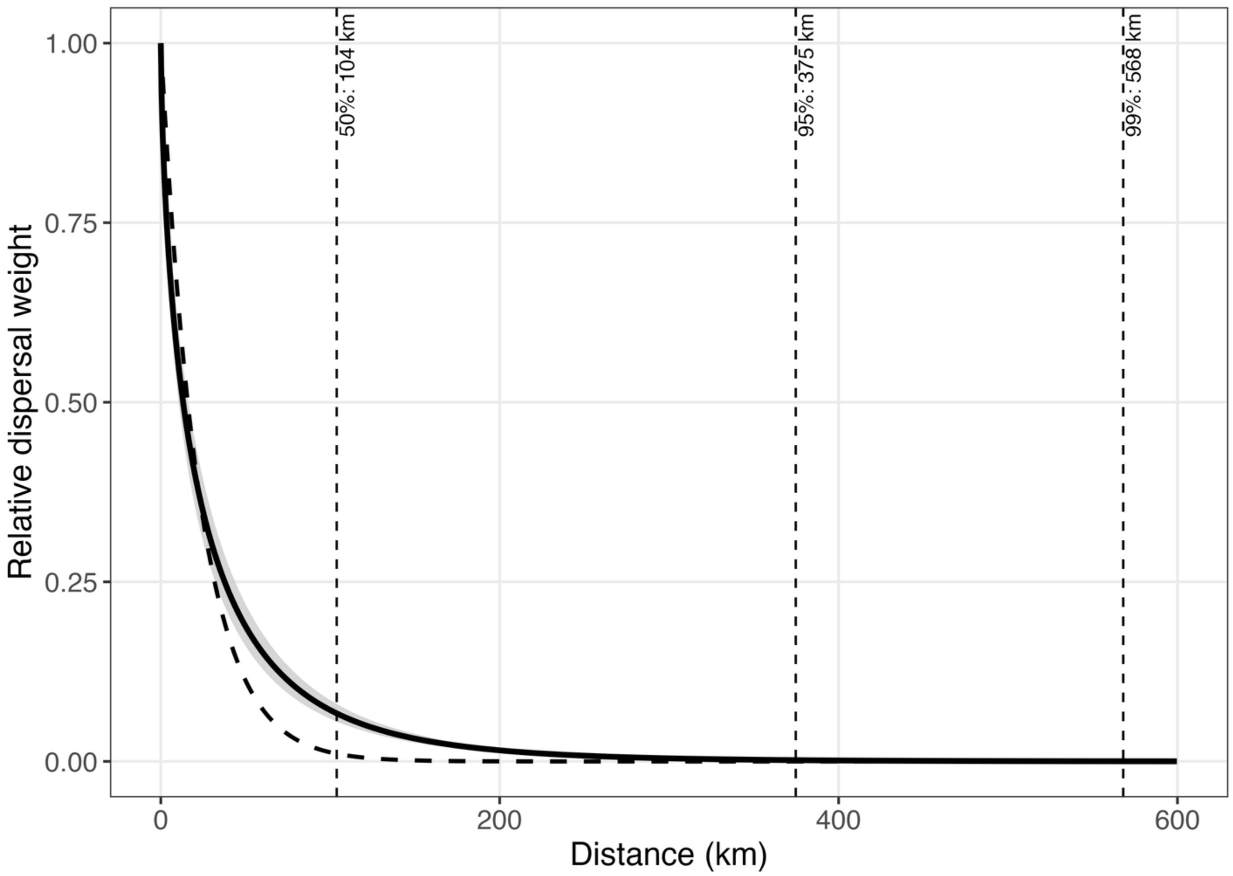


**FIGURE S17.** Distance component of the estimated natal dispersal kernel of German white storks (case study). The solid line shows the posterior mean of $\exp\left(-\left(\frac{d_{i,j}}{\alpha_d}\right)^{\beta_d}\right)$ and the grey ribbon shows the 95% credible interval. Distance is expressed in kilometres between hexagon centroids. The dashed line shows a simple exponential reference with $\beta_d = 1$ and $\alpha_d$ fixed to its posterior median. The vertical lines indicate the posterior mean distances reached by 50%, 95%, and 99% of dispersing storks.

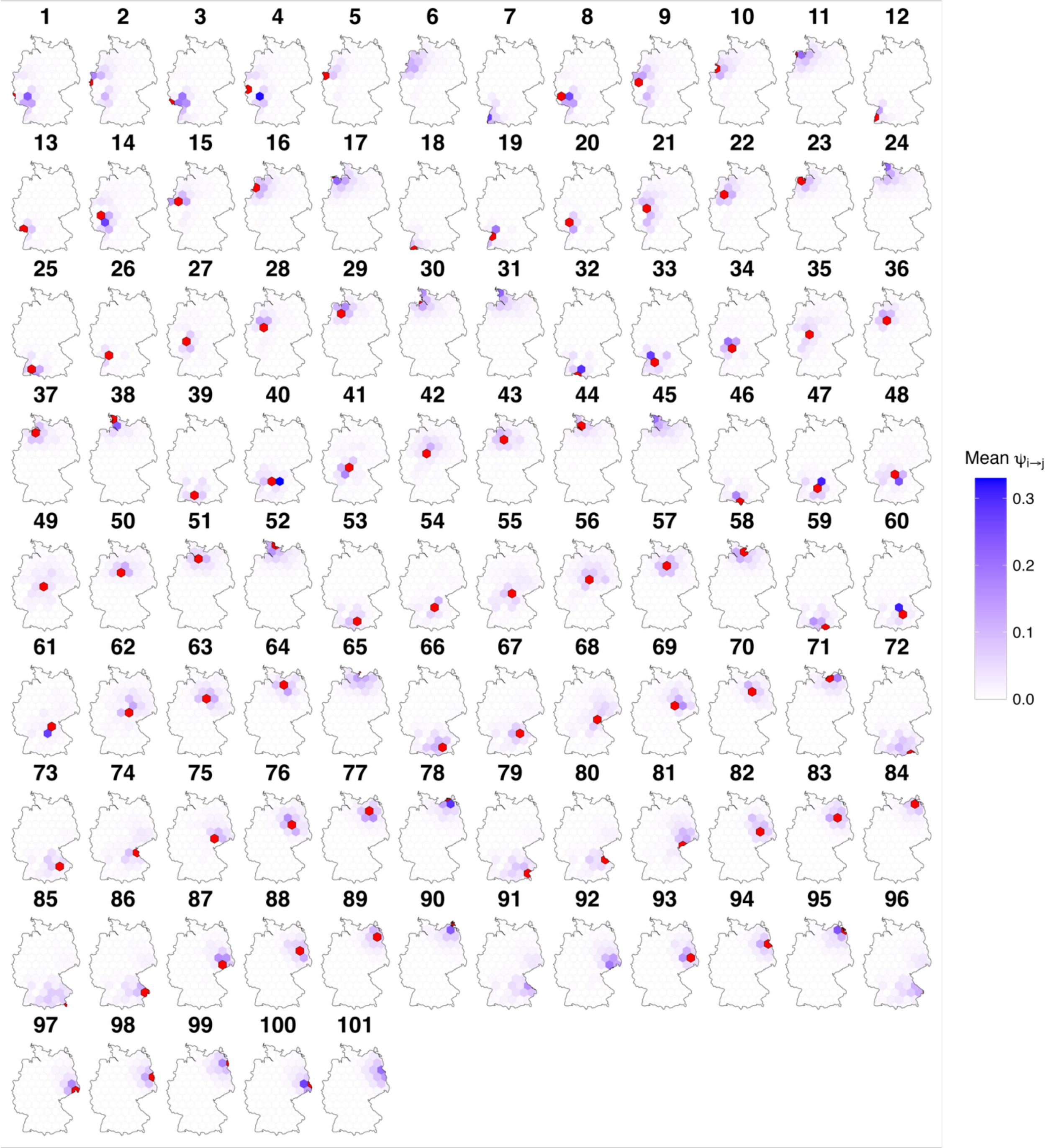


**FIGURE S18.** Posterior mean dispersal probabilities from each of the 101 hexagons. In each panel, the hexagon of origin is highlighted in red.

**TABLE S1.** Notation used in the formulation of the conventional multistate and efficient natal dispersal (END) models.

| Symbol | Definition |
|---|---|
| $T$ | Number of years. |
| $S$ | Number of spatial units. |
| $L$ | Number of rows in the conventional multistate m-array. |
| $A$ | Number of breeder age classes represented in the breeder-END array. |
| $\mathbf{m}$ | Conventional multistate m-array. |
| $\mathbf{m}^{juv}$ | Juvenile-END-array. |
| $\mathbf{m}^{breed}$ | Breeder-END-array. |
| $\boldsymbol{\pi}$ | Cell probabilities of the conventional multistate m-array. |
| $\boldsymbol{\pi}^{juv}$ | Cell probabilities of the juvenile-END-array. |
| $\boldsymbol{\pi}^{breed}$ | Cell probabilities of the breeder-END-array. |
| $R_K$ | Total number of individuals released in state-time combination $K$ (i.e., row of $\mathbf{m}$) in the conventional multistate m-array. |
| $R_{s,t}^{juv}$ | Total number of juveniles released in spatial unit $s$ at time $t$ (i.e., released in a row of the juvenile-END-array). |
| $R_{s,a_{rel},t}^{breed}$ | Total number of breeders released in spatial unit $s$ at time $t$ in breeder age class $a_{rel}$ (i.e., released in a row of the breeder-END-array). |
| $a_{t,u}^{a_{\mathrm{rel}}}$ | Age class in year $u$ of an individual released in age class $a_{\mathrm{rel}}$ in year $t$. |
| $\boldsymbol{\theta}$ | Set of demographic and observation parameters entering the capture-recapture model. |

| Symbol | Definition |
|---|---|
| $\phi_{s,u}^{a}$ | Annual survival probability in spatial unit $s$ at year $u$, for age class $a$. |
| $\gamma_{s,u}^{b}$ | Probability of recruiting in spatial unit $s$ in year $u$ at age $b$, conditional on being alive. |
| $p_{s,u}^{a}$ | Probability that a breeder of age class $a$ is detected in spatial unit $s$ in year $u$. |
| $\psi_{s,r,t}$ | Probability of natal dispersal from spatial unit $s$ to spatial unit $r$ between years $t$ and $t+1$. |
| $\psi_{s,s,t}$ | Fidelity, i.e. the probability of remaining in natal spatial unit $s$ between years $t$ and $t+1$. |
| $q_{s,t,k}$ | Probability that a juvenile settling in spatial unit $s$ after year $t$, and available for detection from year $t+1$ onward, is first detected in year $k$. |
| $g_{s,t,v}$ | Probability of recruiting in year $v$ for an individual that dispersed to spatial unit $s$ between years $t$ and $t+1$ |
| $h_{s,t,k}$ | Probability that an individual that dispersed to spatial unit $s$ from year $t$ to year $t+1$ is first detected in year $k$, after accounting for delayed recruitment |

**TABLE S2.** Predefined distributions used to simulate capture-histories under the different scenarios. For natal dispersal, Dirichlet parameters are ordered here with the fidelity component first for clarity. In the actual transition vector for each natal spatial unit, this value was assigned to the diagonal element, corresponding to movement from a spatial unit to itself, while the remaining values were assigned to movements towards other spatial units.

| **Parameter** | **Scenario(s)** | **Distribution** |
|---|---|---|
| Juvenile survival ($\phi_{juv}$) | 1-10 | $Uniform\ [0.15,0.40]$ |
| 1y survival ($\phi_{1yo}$) | 10 | $Uniform\ [0.60,0.90]$ |
| Adult survival ($\phi_{ad}$) | 1-10 | $Uniform\ [0.75,0.92]$ |
| Detection probability ($p$) | 1-10 | $Uniform\ [0.20,0.90]$ |
| Recruitment at 1y ($\gamma_{1yo}$) | 10 | $Uniform\ [0.10,0.90]$ |
| Natal dispersal ($\psi$) | 1-2, 9-10 | $Dirichlet[3,1,1,1]$ |
| Natal dispersal ($\psi$) | 3-4 | $Dirichlet[9,1,1,1,1,1,1,1,1,1]$ |
| Effect of distance on dispersal ($\alpha_d$) | 5-8 | $Uniform\ [0.8,2.0]$ |

**TABLE S3.** Notation used in the simulation study.

| **Symbol** | **Definition** |
|---|---|
| $S$ | Number of spatial units (either 4 or 10, depending on the scenario). |
| $N$ | Number of simulation replicates (100). |
| $\psi_{i,j}$ | Natal dispersal probability from an origin spatial unit $i$ to a destination spatial unit $j$. |

| **Symbol** | **Definition** |
|---|---|
| $d_{i,j}$ | Euclidean distance between the centroids of an origin ($i$) and a destination ($j$) spatial unit. |
| $\alpha_d$ | Parameter controlling the effect of distance on natal dispersal probability in the distance-based simulation scenarios. |
| $\hat{\theta}_n$ | Posterior mean of the parameter $\theta$ in a given simulation replicate. |
| $\theta_n$ | True value of the parameter $\theta$ in a given simulation replicate. |
| $\text{RMSE}_\theta$ | Root mean squared error of the parameter $\theta$ across simulation replicates. |
| $\text{Coverage}_\theta$ | Proportion of simulation replicates for which the 95% credible interval contains the true parameter ($\theta$) value. |
| $L_n^\theta$ | Lower bound of the 95% credible interval of parameter $\theta$ for simulation replicate $n$. |
| $U_n^\theta$ | Upper bound of the 95% credible interval of parameter $\theta$ for simulation replicate $n$. |

**TABLE S4.** Notation used in the White Stork case study.

| Symbol | Definition |
|---|---|
| $H$ | Number of hexagonal spatial units in the case study (101). |
| $\mu_i^\theta$ | Hexagon-specific mean, on the logit scale, of the temporally varying parameter $\theta$. |
| $\sigma^\theta$ | Standard deviation of the temporal random effect of the temporally varying parameter $\theta$. |

| Symbol | Definition |
| --- | --- |
| $\mu^{\zeta}$ | Overall mean of the spatially varying parameter $\zeta$. |
| $\tau^{\zeta}$ | Precision parameter of the ICAR distribution, controlling the strength of spatial autocorrelation in the spatially varying parameter $\zeta$. |
| $\phi_{i,t}^{juv}$ | Annual juvenile survival probability in a given hexagon $i$ and year $t$. |
| $\phi_{i,t}^{ad}$ | Annual adult survival probability in a given hexagon $i$ and year $t$. |
| $\gamma_i^b$ | Probability of recruiting at age $b$ in hexagon $i$, conditional on being alive; recruitment is temporally constant in the case study. |
| $p_{x,i,t}$ | Adult breeder resighting probability in a given hexagon $i$ and year $t$, conditional on previous-year resighting status; the $x$ index equals 1 for individuals resighted in the previous year and 2 otherwise. |
| $\psi_{i,j}$ | Probability of natal dispersal from an origin hexagon $i$ to a destination hexagon $j$. |
| $\alpha_d$ | Scale parameter of the power-exponential natal dispersal kernel. |
| $\beta_d$ | Shape parameter of the power-exponential natal dispersal kernel. |
| $d_{i,j}$ | Distance, standardized between 0 and 1, between the centroids of an origin ($i$) and a destination ($j$) hexagon. |
| $P_j$ | Mean population size in the destination hexagon $j$ across the study period. |
| $\gamma_{pop}$ | Effect of destination population size on natal dispersal probability. |
| $\eta_{i,j}$ | Residual origin ($i$)-destination ($j$) effect on natal dispersal not explained by distance or destination population size. |
| $\sigma^{\eta}$ | Standard deviation of the residual origin-destination effects. |
| $\vartheta_i$ | Set of hexagons neighbouring a focal hexagon $i$ in the ICAR formulation. |

| Symbol | Definition |
| --- | --- |
| $V_i$ | Number of hexagons neighbouring a focal hexagon $i$ in the ICAR formulation. |
| $\alpha_i^b$ | Probability of first breeding at a given age $b$ in hexagon $i$. |
| $M_i$ | Mean age at first breeding in hexagon $i$. |
| $\mu_M$ | Overall mean age at first breeding. |
| $\mathbf{J}_{i,t}$ | Vector containing the numbers of juveniles produced in an origin hexagon $i$ at year $t$ that survive and settle in each destination hexagon, together with the number that die (last component). |
| $F_{i,t}$ | Number of juveniles produced in hexagon $i$ at year $t$. |
| $\boldsymbol{\delta}_{i,t}$ | Vector containing the probabilities that juveniles produced in an origin hexagon $i$ at year $t$ survive and settle in each destination hexagon, together with the probability that they die (last component); cell probabilities of $\mathbf{J}_{i,t}$. |
| $I_{i,t}$ | Number of juvenile immigrants settling in a hexagon $i$ at year $t$. |
| $E_{i,t}$ | Number of juvenile emigrants leaving hexagon $i$ (i.e., settling in another hexagon) at year $t$. |
| $I_{i,t}^{rec}$ | Expected number of immigrants recruiting into hexagon $i$ at year $t$. |
| $E_{i,t}^{rec}$ | Expected number of individuals originating from hexagon $i$ and recruiting elsewhere in year $t$. |
| $\lambda_{i,t}$ | Observed population growth rate between two consecutive years ($t-1$ and $t$) in hexagon $i$. |
| $C_{i,t}$ | Intrinsic growth rate in hexagon $i$ and year $t$, corrected for recruiting immigration and emigration. |
| $\bar{C}_i$ | Geometric mean intrinsic growth rate for hexagon $i$ over 2005-2023. |

| Symbol | Definition |
| --- | --- |
| $G_{i,t}$ | Number of breeding individuals in hexagon $i$ and year $t$. |
| $B_{i,t}$ | Total number of breeding pairs in hexagon $i$ and year $t$. |
| $B_{i,t}^{obs}$ | Number of breeding pairs for which juvenile production was observed in hexagon $i$ and year $t$. |
| $B_{i,t}^{mis}$ | Number of breeding pairs for which juvenile production was missing in hexagon $i$ and year $t$. |
| $f_{i,t}$ | Productivity, defined as the expected number of fledged juveniles per breeding pair in hexagon $i$ and year $t$. |
| $F_{i,t}^{obs}$ | Number of fledged juveniles observed among sampled breeding pairs in hexagon $i$ and year $t$. |
| $F_{i,t}^{mis}$ | Estimated number of fledged juveniles produced by non-sampled breeding pairs in hexagon $i$ and year $t$. |
| $D_{i,j,t}$ | Expected number of juveniles fledged in an origin hexagon $i$ in a given year $t$ that survive and settle in a destination hexagon $j$. |
| $O_{i,j,t}$ | Expected number of individuals produced in an origin hexagon $i$ and recruiting in a destination hexagon $j$ in year $t$. |

**TABLE S5.** Priors assigned to stochastic parameters in the White stork case study.

| **Parameter** | **Prior** |
| --- | --- |
| Overall mean juvenile survival (at logit-scale; $\mu_{juv}^{\phi}$) | $Normal\ (\mu = 0, sd = 3)$ |
| Overall mean adult survival (at logit-scale; $\mu_{ad}^{\phi}$) | $Normal\ (\mu = 0, sd = 3)$ |

| | |
|---|---|
| Overall mean resighting probability if seen the year before (at logit-scale; $\mu^{p}_{x=1}$) | $Normal\ (\mu = 0, sd = 3)$ |
| Overall mean resighting probability if not seen the year before (at logit-scale; $\mu^{p}_{x=2}$) | $Normal\ (\mu = 0, sd = 3)$ |
| Overall, age-dependent mean recruitment probability ($\mu^{\gamma}_{a}$) | $Normal\ (\mu = 0, sd = 10)$ |
| Precision parameter of the ICAR spatial random effect for juvenile survival ($\tau^{\phi}_{juv}$) | $Gamma\ (\alpha = 0.001, \beta = 0.001)$ |
| Precision parameter of the ICAR spatial random effect for adult survival ($\tau^{\phi}_{ad}$) | $Gamma\ (\alpha = 0.001, \beta = 0.001)$ |
| Precision parameter of the ICAR spatial random effect for resighting probability if seen the year before ($\tau^{p}_{x=1}$) | $Gamma\ (\alpha = 0.001, \beta = 0.001)$ |
| Precision parameter of the ICAR spatial random effect for resighting probability if not seen the year before ($\tau^{p}_{x=2}$) | $Gamma\ (\alpha = 0.001, \beta = 0.001)$ |
| Precision parameter of the ICAR spatial random effect for recruitment ($\tau^{\gamma}_{a}$) | $Gamma\ (\alpha = 0.001, \beta = 0.001)$ |
| Temporal standard deviation of juvenile survival ($\sigma^{\phi}_{juv}$) | $Uniform\ (0, 10)$ |
| Temporal standard deviation of adult survival ($\sigma^{\phi}_{ad}$) | $Uniform\ (0, 10)$ |
| Temporal standard deviation of resighting probability if seen the year before ($\sigma^{p}_{x=1}$) | $Uniform\ (0, 10)$ |

| | |
|---|---|
| Temporal standard deviation of resighting probability if not seen the year before ($\sigma^{p}_{x=2}$) | $Uniform\ (0, 10)$ |
| Distance-scale parameter of dispersal ($\alpha_d$) | $Uniform\ (0, 10)$ |
| Distance-decay shape parameter of natal dispersal ($\beta_d$) | $Uniform\ (0.5, 3)$ |
| Effect of destination population size on dispersal ($\gamma_{pop}$) | $Normal\ (\mu = 0, sd = 2)$ |
| Standard deviation of residual variation in natal dispersal ($\sigma^{\eta}$) | $Uniform\ (0, 5)$ |
