## Appendix S1 for "Efficient capture–recapture inference for spatially varying natal dispersal, survival and recruitment"

**APPENDIX S1. MODELLING DETAILS**

In this appendix we describe some modelling details, in the following order:

1. Computation of $g_{s,t,v}$ (END model)

2. Calculation of root mean squared error and coverage (simulation study)

3. ICAR formulation (case study)

4. Calculation of mean population size per hexagon (case study)

5. Test 2.CT for assessment of immediate trap-response (case study)

6. Calculation of age at first breeding (case study)

7. Calculation of the number of juveniles produced per hexagon (case study)

8. Calculation of expected numbers of immigrants and emigrants (source-sink dynamics; case study)

### 1. Computation of $g_{s,t,v}$ (END model)

For an individual that dispersed to spatial unit $s$ between years $t$ and $t+1$, the probability $g_{s,t,v}$ of recruiting in year $v$ is

$$g_{s,t,v} = \begin{cases} \gamma_{s,t+1}^{b_{t,t+1}}, & if\ v = t+1, \\ \gamma_{s,v}^{b_{t,v}} \prod_{u=t+1}^{v-1} \left[ \left(1-\gamma_{s,u}^{b_{t,u}}\right) \phi_{s,u}^{a_{t,u}} \right], & if\ v > t+1, \end{cases}$$

where $\gamma_{s,v}^{b_{t,v}}$ is the probability of recruiting in spatial unit $s$ at time $v$ and for age $b_{t,v}$, conditional on being alive. If recruitment is certain by a maximum age (as generally assumed), the recruitment probability at that age is fixed to 1 for individuals that have not recruited previously.

For example, when some individuals recruit at age 1 with probability $\gamma_{s,v}^{1}$, and the others recruit at age 2, given that they have survived until age 2, then

$$g_{s,t,t+1} = \gamma_{s,t+1}^{1},$$

and

$$g_{s,t,t+2} = \left(1-\gamma_{s,t+1}^{1}\right) \phi_{s,t+1}^{1yo},$$

where $\phi_{s,t+1}^{1yo}$ is survival of 1-year-old individuals in spatial unit $s$ at time $t+1$.

### 2. Calculation of root mean squared error and coverage (simulation study)

We calculated the root mean squared error (RMSE) for each parameter $\theta$ as

$$RMSE_\theta = \sqrt{\frac{1}{N}\sum_{n=1}^{N}(\widehat{\theta_n} - \theta_n)^2},$$

where $N$ is the number of simulation replicates (here always 100), $\widehat{\theta_n}$ is the posterior mean of the parameter in the model for simulation $n$, and $\theta_n$ the true value of the parameter in simulation $n$.

We calculated the coverage as

$$Coverage_\theta = \frac{1}{N}\sum_{n=1}^{N} I\left(L_n^\theta \le \theta_n \le U_n^\theta\right),$$

where $L_n^\theta$ and $U_n^\theta$ are the lower and upper bounds of the 95% credible interval of parameter $\theta$ for simulation $n$, and $I(\cdot)$ is an indicator function equal to 1 when the interval contained the true value and 0 otherwise.

### 3. ICAR formulation (case study)

An intrinsic conditional autoregressive (ICAR; Besag, 1974) model was used to model spatial autocorrelation among neighbouring hexagons in survival, recruitment and resighting probability. The rationale of this approach is that the parameter value estimated for a given hexagon is informed by the parameter values of neighbouring hexagons, such that neighbouring hexagons tend to have more similar values than non-neighbouring hexagons. In our case, a neighbouring hexagon to hexagon $i$ was simply defined as a hexagon sharing a border with $i$.

For any spatially correlated parameter $\zeta$, we denote the vector of spatially varying values on the linear predictor scale as

$$\boldsymbol{\zeta} = (\zeta_1, \dots, \zeta_H),$$

where each element $\zeta_i$ corresponds to the value of the parameter in hexagon $i$. Under the ICAR prior, the full conditional distribution of $\zeta_i$, given all other values $\zeta_{-i}$, is

$$\zeta_i \mid \zeta_{-i} \sim Normal\left(\frac{1}{V_i}\sum_{g\in\vartheta_i} \zeta_g, \frac{1}{V_i \tau^{\zeta}}\right),$$

where $\vartheta_i$ is the set of neighbouring hexagons of hexagon $i$, $V_i = |\vartheta_i|$ is the number of neighbours of hexagon $i$, and $\tau^{\zeta}$ is the precision parameter of the ICAR distribution. This precision parameter determines the amount of spatial smoothing: larger values of $\tau^{\zeta}$ imply lower variance around the neighbourhood mean, and therefore stronger similarity among neighbouring hexagons. Each parameter $\zeta$ was assigned its own precision parameter, allowing the strength of spatial smoothing to differ among parameters.

The ICAR formulation centres each parameter value on the average value of its neighbours, thereby inducing positive spatial autocorrelation. Because the intrinsic CAR prior is improper, its precision matrix is singular. To ensure identifiability, we imposed the standard sum-to-zero constraint on deviations from the overall mean,

$$\sum_i (\zeta_i - \mu^{\zeta}) = 0.$$

### 4. Calculation of mean population size per hexagon (case study)

Mean population size was calculated from annual population data available at the Kreis (municipal) level from 2000 to 2022. In most cases, these data were available for individual Kreise, although some Kreise were occasionally grouped (see provided data and Supplement: Fig. S15). To transfer population size from Kreise to hexagons, we first calculated the proportion of each Kreis overlapping with each hexagon. Because Kreise were generally smaller than the hexagons, many Kreise were entirely contained within a single hexagon. When a Kreis overlapped more than one hexagon, its population size was allocated proportionally to the area of the Kreis falling within each hexagon. For example, if a Kreis contained two breeding pairs and 50% of its area overlapped with a given hexagon, one breeding pair was assigned to that hexagon. This approach assumes that population size was uniformly distributed within each Kreis.

For each year, population size in each hexagon was then obtained by summing the area-weighted contributions of all Kreise overlapping that hexagon. Finally, we calculated the mean population size of each hexagon across the full study period. If population size was unavailable for a hexagon in a given year because at least one contributing Kreis had missing data, that year was excluded from the calculation of the mean. This occurred rarely: no hexagon had more than one missing year, and only four hexagons had any missing annual value.

### 5. Test 2.CT for assessment of immediate trap-response (case study)

To prevent spatial heterogeneity in resighting probability from being interpreted as immediate trap-response, adult breeder encounter histories were analysed separately for each hexagon. When an individual was observed in multiple hexagons, only its resightings within the focal hexagon were retained for the analysis of that hexagon. At each year, Test 2.CT compares the probability of being resighted at the next occasion between individuals that were and were not resighted at the current occasion. R2Ucare calculated a test statistic and its degrees of freedom for every year in which this comparison could be performed, and then summed them across years to obtain the hexagon-specific result of Test 2.CT. To obtain the overall test, we summed the test statistics and degrees of freedom across hexagons and calculated the p-value from a chi-squared distribution with the resulting degrees of freedom. To determine the direction of the response, we summed the yearly signed statistics across all hexagons and divided their sum by the square root of their total number, following the procedure used by Test 2.CT to combine years. The overall stratified test indicated trap-happiness ($\chi^2$ = 2416.20, df = 731, P < 0.001; signed statistic = −17.77).

### 6. Calculation of age at first breeding (case study)

Age at first breeding was derived from the age-specific recruitment probabilities estimated by the model. Recruitment probability at age $b$ in hexagon $i$, denoted $\gamma_i^b$, was defined in the model conditionally on an individual not having recruited at any younger age. We therefore converted these conditional recruitment probabilities into absolute probabilities of first breeding at each age. Thus, the probability of first breeding at age 1 was $\gamma_i^1$, whereas the probability of first breeding at age 2 was $\gamma_i^2(1-\gamma_i^1)$. The same sequential calculation was applied up to age 4, and all remaining individuals were assumed to recruit at age 5.

For each hexagon, the age at first breeding $M_i$ was then calculated as the weighted average of ages 1 to 5, using the absolute probabilities of first breeding at each age as weights:

$$M_i = \sum_{b=1}^{5} b\alpha_i^b,$$

where $\alpha_i^b$ is the absolute probability of first breeding at age $b$ in hexagon $i$.

The same procedure was used to calculate the overall mean age at first breeding $\mu^M$, using in this case the overall recruitment probabilities.

### 7. Calculation of the number of juveniles produced per hexagon (case study)

Data on juvenile production was not sampled in all Kreise and years. We therefore first identified, for each Kreis and year, the number of fledged juveniles that had been counted, together with the number of breeding pairs for which juvenile production was known. These data were transferred to hexagons using the area-weighted allocation described in section 3, and the resulting values were rounded to the nearest integer. This provided, for each hexagon and year, the number of fledged juveniles observed in the sampled part of the population and the corresponding number of sampled breeding pairs. In parallel, we also calculated the total number of breeding pairs per hexagon and year using the same area-weighted allocation. When the number of breeding pairs was missing for a single year in a

Kreis, the missing value was replaced by the mean of the two previous and the two following years, rounded to the nearest integer.

For hexagon-year combinations with complete juvenile data (i.e., when the number of sampled breeding pairs was equal to the total number of breeding pairs), the observed number of juveniles was retained unchanged. For incomplete hexagon-year combinations, the total number of juveniles was estimated using a separate model fitted in NIMBLE. Let $B_{i,t}$ be the total number of breeding pairs in hexagon $i$ and year $t$, $B_{i,t}^{obs}$ the number of breeding pairs for which the number of juveniles produced was known, and $B_{i,t}^{mis} = B_{i,t} - B_{i,t}^{obs}$ the number of breeding pairs for which data on juveniles are missing. For each hexagon-year combination with incomplete juvenile data, we estimated the productivity $f_{i,t}$, defined as the expected number of fledged juveniles per breeding pair, from the number of sampled juveniles $F_{i,t}^{obs}$:

$$F_{i,t}^{obs} \sim Poisson\ (f_{i,t} B_{i,t}^{obs}).$$

The same productivity was then used to estimate the number of juveniles produced by non-sampled breeding pairs:

$$F_{i,t}^{mis} \sim Poisson(f_{i,t} B_{i,t}^{mis}).$$

The total number of juveniles produced in incomplete hexagon-year combinations was then computed as

$$F_{i,t} = F_{i,t}^{obs} + F_{i,t}^{mis}.$$

In the model, we assigned each $f_{i,t}$ an independent $Uniform\ (0.5, 2.5)$ prior. The juvenile production model used the same MCMC settings as the END model: four chains of 40,000 iterations, of which 5,000 were discarded as burn-in, with a thinning interval of 20. For each of the 1,750 retained draws per chain, we constructed a complete matrix of the number of juveniles produced per year and hexagon by retaining the observed values for complete hexagon-year combinations and using the corresponding posterior draw for incomplete combinations. These matrices represented the posterior distribution of juvenile production across hexagons and years.

#### 8. Calculation of expected numbers of immigrants and emigrants (source-sink dynamics; case study)

To calculate $I_{i,t}^{rec}$ and $E_{i,t}^{rec}$, we first derived the expected number of juveniles fledged in hexagon $i$ in year $t$ that survived and settled in hexagon $j$:

$$D_{i,j,t} = F_{i,t}\ \phi_{i,t}^{juv}\ \psi_{i,j}.$$

We then converted these expected dispersed juveniles into expected numbers of recruits. For a juvenile that settled in hexagon $j$ after dispersal in year $t$, the expected contribution to recruitment at age $b$ was calculated as the probability of not recruiting before age $b$, surviving until that age, and recruiting at age $b$. These expected recruits were assigned to year $t + b$, i.e. the year of recruitment rather than the year of natal dispersal. For example, juveniles moving from $i$ to $j$ in year $t$ contributed

$$D_{i,j,t}\gamma_j^1$$

expected recruits in hexagon $j$ in year $t+1$, and

$$D_{i,j,t}\left(1-\gamma_j^1\right)\phi_{j,t+1}^{ad}\gamma_j^2$$

expected recruits in hexagon $j$ in year $t+2$. The same calculation was extended up to age 5, the maximum recruitment age considered in the model.

Let $O_{i,j,t}$ denote the expected number of individuals produced in hexagon $i$ and recruiting in hexagon $j$ in year $t$, after summing over all relevant natal years. The expected number of immigrants recruiting into hexagon $i$ in year $t$ was then calculated as

$$I_{i,t}^{rec} = \sum_{j \neq i} O_{j,i,t},$$

whereas the expected number of emigrants from hexagon $i$ recruiting elsewhere in year $t$ was calculated as

$$E_{i,t}^{rec} = \sum_{j \neq i} O_{i,j,t}\,.$$
